# The Dsc ubiquitin ligase complex identifies transmembrane degrons to degrade orphaned proteins at the Golgi

**DOI:** 10.1101/2024.03.11.584465

**Authors:** Yannick Weyer, Sinead Iduna Schwabl, Xuechen Tang, Michael A. Widerin, Veronika Niedrist, Maria G. Tettamanti, Sabine Weys, Bettina Sarg, Leopold Kremser, Klaus R. Liedl, Oliver Schmidt, David Teis

**Author notes:** Yannick Weyer and Sinead Iduna Schwabl contributed equally to this work. corresponding author: David Teis, PhD Institute of Molecular Biochemistry Medical University of Innsbruck Innrain 80/82 CCB Building A-6020, Innsbruck, Austria.

## Abstract

The Golgi apparatus is essential for protein sorting, yet its quality control mechanisms are poorly understood. Here we show that the Dsc ubiquitin ligase complex, particularly the rhomboid pseudo-protease subunit, Dsc2, assesses the hydrophobic length of α-helical transmembrane domains (TMDs) at the Golgi. Thereby the Dsc complex interacts with orphaned ER and Golgi proteins that have shorter TMDs and ubiquitinates them for targeted degradation. Some Dsc substrates will be K63 polyubiquitinated for ESCRT dependent vacuolar degradation or K48 polyubiquitinated for *e*ndosome and *G*olgi *a*ssociated proteasomal *d*egradation (EGAD). Other Dsc substrates are exclusively extracted by Cdc48 for EGAD. The accumulation of Dsc substrates entails a specific increase in glycerophospholipids with shorter and asymmetric fatty acyl chains. Hence, the Dsc complex mediates the selective degradation of orphaned proteins at the sorting center of cells, which prevents their spreading across other organelles and thus preserves cellular membrane protein and lipid composition.

## Introduction

The Golgi apparatus is essential for the sorting of all proteins from the endoplasmic reticulum (ER) to the endo-lysosomal system, or along the secretory pathway to the PM or to the extracellular space ^1^ ^2^ ^3^. Despite the pivotal role of the Golgi in protein sorting, and in cellular - and organismal health, Golgi-based protein quality control (PQC) processes have remained largely unknown. While there is emerging evidence that PQC systems operate at the Golgi ^4-7^ to detect and degrade misfolded and orphaned proteins (proteins that mislocalize or fail to assemble with their protein binding partners ^8,9^), the underlying molecular mechanisms are not well characterized ^10-12^.

A candidate for detecting and degrading orphaned membrane proteins at the Golgi in yeast is the ‘Defective in SREBP cleavage’ Dsc E3 ubiquitin ligase complex ^7,13^. The Dsc complex is a multi-subunit transmembrane protein complex, that localizes to the Golgi, endosomes and to the limiting membrane of the vacuole. The RING E3 ligase Tul1 forms a complex with Dsc2, Dsc3, Ubx3 ^13,14^, and with one of two trafficking adaptors, Gld1 or Vld1 ^15^. Gld1 targets the Dsc complex to Golgi and endosomes. Vld1 diverts the Dsc complex from the Golgi directly to the limiting membrane of the vacuole via the AP3 pathway. Tul1 contains at its N-terminus a large luminal domain, seven predicted TMDs and a C-terminal cytosolic RING E3 ligase domain, which interacts with the E2 ubiquitin conjugating enzyme Ubc4 ^16^. Tul1 is conserved in a wide range of fungi and plants ^7,17^. Direct human orthologues are not obvious ^7^. Dsc2 is a rhomboid like pseudo-protease, with similarities to Derlins (Der1, Dfm1) and UBA domain containing 2 (UBAC2). Dsc3 contains a Ubl domain, and shares some similarities to the transmembrane and ubiquitin-like (Ubl) domain containing protein 1/2 (TMUB1/2). Ubx3 has a C-terminal UBX domain that can bind to Cdc48 ^18^, and is related to the FAS-associated factor 2 (FAF2/UBXD8). Overall, the subunit architecture of the Dsc complex resembles ubiquitin ligase complexes that function in ER associated degradation (ERAD) ^19-22^.

The Dsc complex can ubiquitinate ectopically expressed, non-native proteins at Golgi and endosomes, ^16,23,24^. The Vld1-Dsc complex ubiquitinates native vacuolar membrane proteins (mainly lysosomal transmembrane transporters) ^25,26^. Once these proteins are ubiquitinated, they are recognized by the *e*ndosomal *s*orting *c*omplexes *r*equired for *t*ransport (ESCRT) and targeted into vacuoles for degradation. The Dsc complex also has the capacity to ubiquitinate phosphatidylethanolamine (PE), which helps to recruit the ESCRT machinery ^27^.

Recently, we demonstrated that the Gld1-Dsc complex polyubiquitinates the ER resident membrane protein Orm2 on two cytosolic lysine residues (K25, K33) for Cdc48-mediated membrane extraction and proteasomal degradation, as soon as Orm2 exits the ER and enters the Golgi. We termed this membrane protein degradation pathway *e*ndosome and *G*olgi *a*ssociated *d*egradation (EGAD) ^7,20,28^. To reach the Golgi, Orm2 must no longer bind to the serine palmitoyl transferase complex (SPT, Lcb1/2) at the ER, and exit the ER in COPII carriers with the help of the derlin Dfm1 ^29^. EGAD of Orm2 contributes to the de-repression of SPT activity and thus helps to maintain sphingolipid homeostasis ^7^. Besides Orm2, the native substrate spectrum of the Gld1-Dsc complex was unknown. It was also unclear how the Dsc complex detected its substrates, and which substrates were degraded via EGAD or ESCRT pathways.

Now we have used a combination of quantitative proteomics and cell biological approaches to (i) identify additional substrates of the Dsc complex (ii), define the biochemical properties of their TMDs, (iii) reveal how they are detected and degraded and (v) determined the cellular consequences, when Dsc substrates accumulate. Our results suggest that the Dsc complex uses Dsc2 to detect the hydrophobic length of TMDs and thereby selectively degrades orphaned ER and Golgi membrane proteins that expose short TMDs at the Golgi. Some Dsc substrates (Orm2, Yif1) are degraded exclusively by EGAD, while others (e.g.: the heme oxygenase Hmx1) can be degraded either via EGAD - or ESCRT pathways. The accumulation of Dsc substrates (membrane proteins with short TMDs) causes an unusual change of the cellular lipid composition, with an increase of glycerophospholipid species with shorter and asymmetric acyl chains. Thus, the Gld1-Dsc complex restricts the accumulation of orphaned proteins at the Golgi, and prevents them from spreading to endosomes and to the limiting membrane of vacuoles. This Golgi quality control is essential to maintain cellular membrane protein and lipid composition.

## Results

### Dsc complex substrates differ from ESCRT clients

To understand how the Dsc ubiquitin ligase complex maintains membrane homeostasis, we compared the membrane protein and lipid composition of cells lacking the E3 ligase Tul1 (*tul1Δ*) to isogenic WT cells.

First, we used stable isotope labeling with amino acids in cell culture (SILAC) to identify changes in the membrane proteome of *tul1Δ* cells relative to WT cells (labeled with “heavy” [^13^C_6_,^15^N_2_]-L-lysine). We quantified 667 membrane proteins (52% of all membrane proteins), out of 3631 quantified proteins (n=3, approximately 90% of the yeast proteome) (Figure 1A,B, Table S1,S2) ^8,30-33^. In the *tul1Δ* cells, 21 membrane proteins were upregulated (3% of the quantified membrane proteins) (Figure 1B, Table S2). Many upregulated proteins were ER and Golgi proteins (Alg3, Csg2, Dgk1, Elo2, Hmx1, Orm2, Tpo5, Neo1, Gnt1) (Figure 1B, Table S3). The bona fide Dsc substrate Orm2 was among the upregulated proteins ^7^. Additionally, the heme oxygenase Hmx1, the phospholipid flippase Neo1, and the high affinity iron permease Ftr1 were identified in our earlier work as potential Dsc complex substrates ^7^.

**Figure 1:**
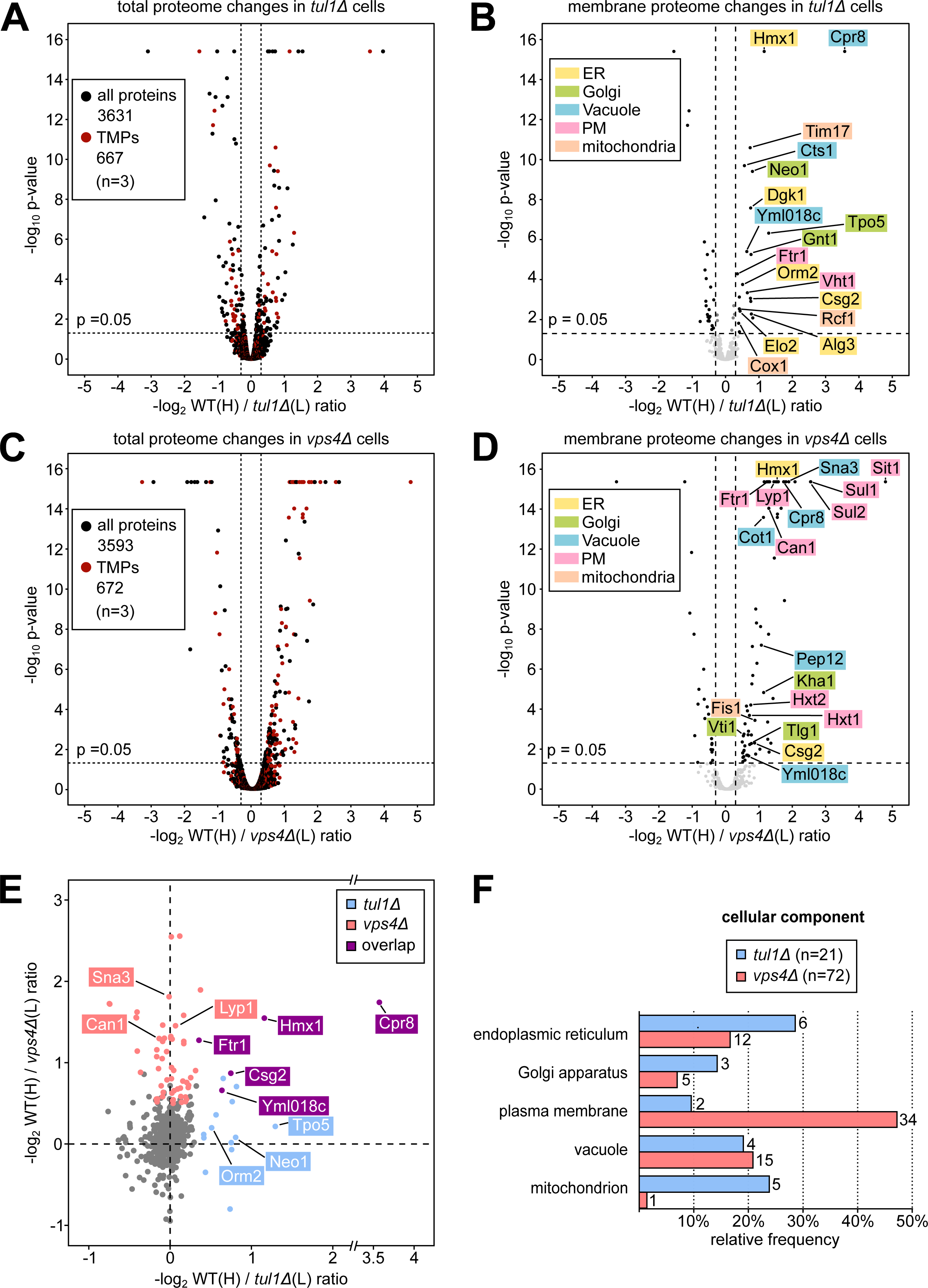
Quantitative proteomic SILAC profiling of Dsc complex and ESCRT mutants. **(A-D)** Volcano plots showing the H/L peptide ratios of proteins from heavy ^13^C_6_,^15^N_2_-L-lysine labeled WT cells against light ^12^C_6_,^14^N_2_-L-lysine L-lysine labeled *tul1Δ* **(A, B)** or *vps4Δ* **(A, D)** mutants. Total proteome of **(A)** WT / *tul1Δ* mutants or **(C)** WT / *vps4Δ* mutants, transmembrane proteins (TMPs) in red; **(B, D)** only transmembrane proteome. Significantly regulated (-log_2_ ≤ 0.3, ≥ 0.3, p ≤ 0.05) transmembrane membrane proteins in black. See also **Tables S1, S2**. **(E)** Scatter plot of the 614 membrane proteins quantified in WT/*tul1Δ* (x-axis) and in WT/*vps4Δ* (y-axis), only displaying proteins with -log>-1. Proteins are significantly upregulated (-log_2_ ≥ 0.3, p ≤ 0.05) in both datasets are highlighted: Dsc complex-dependent (light blue), ESCRT-dependent (light red) and overlapping substrates (purple). See also **Table S4**. **(F)** Gene Ontology (GO) analysis for cellular components of the significantly upregulated transmembrane proteins in *tul1Δ* (light blue) or *vps4Δ* (light red) cells. Only GO terms with ≥ 2 hits are shown as relative frequency over the data set. See also **Table S3.**

To understand how the potential Dsc substrates differed from ESCRT clients, we determined how loss of ESCRT function in *vps4Δ* mutant cells changed the membrane proteome (Figure 1C, D Table S1, S2). In total, we quantified 672 membrane proteins in the WT/*vps4Δ* cell extracts (Figure 1D, Table S2). Of those 90% (614) were also quantified in WT/*tul1Δ* cell extracts (Figure 1E, Table S4). In ESCRT mutants 72 membrane proteins were upregulated (> 10% of the quantified membrane proteins), of which many (34) were plasma membrane proteins (Figure 1D&E, Table S3), including well known clients of the ESCRT machinery (e.g.: Can1, Lyp1). Most proteins that were upregulated in *vps4Δ* cells, were not upregulated in *tul1Δ* cells (and vice versa) (Figure 1E). Five proteins (Hmx1, Crp8, Csg2, Ftr1, Yml018c) were significantly upregulated in both *tul1Δ* and *vps4Δ* mutants, suggesting that these could be substrates of the Dsc complex, which are targeted to the ESCRT pathway for vacuolar degradation.

Our census of the membrane protein composition showed that the ESCRT machinery and the Dsc complex played largely complementary roles in controlling the cellular membrane protein composition.

### Dsc substrates are ER and Golgi proteins with short α-helical transmembrane domains

To identify characteristic signatures of Dsc substrates, we compared the α-helical transmembrane domains (TMDs) of the proteins that were upregulated in *tul1Δ* and *vps4Δ* mutants. TMDs were predicted using DeepTMHMM ^34^. For the 21 membrane proteins upregulated in *tul1*Δ cells, DeepTMHMM predicted 125 TMDs in 18 proteins. For the 72 proteins up-regulated in *vps4*Δ cells, 550 TMDs were predicted in 64 proteins (Figure 2A, Table S5). In *tul1*Δ cells, proteins with relatively short TMDs (hydrophobic length) accumulated: 44% of the TMDs contained ≤ 18 amino acids (Figure 2A, i). 66% of the proteins had at least one TMD with ≤ 16 amino acids, and every protein had at least one TMD with ≤18 amino acids (Figure 2A, ii), with 75% of these proteins having ≤ 10 TMDs (Figure 2A, iii). In *vps4*Δ cells, the accumulating proteins showed a preference for longer TMDs and more TMDs per protein: Only 26% of the TMDs were short with ≤ 18 amino acids (Figure 2A, i – ii) with 48% of these proteins had > 10 TMDs (Figure 2A, iii).

**Figure 2:**
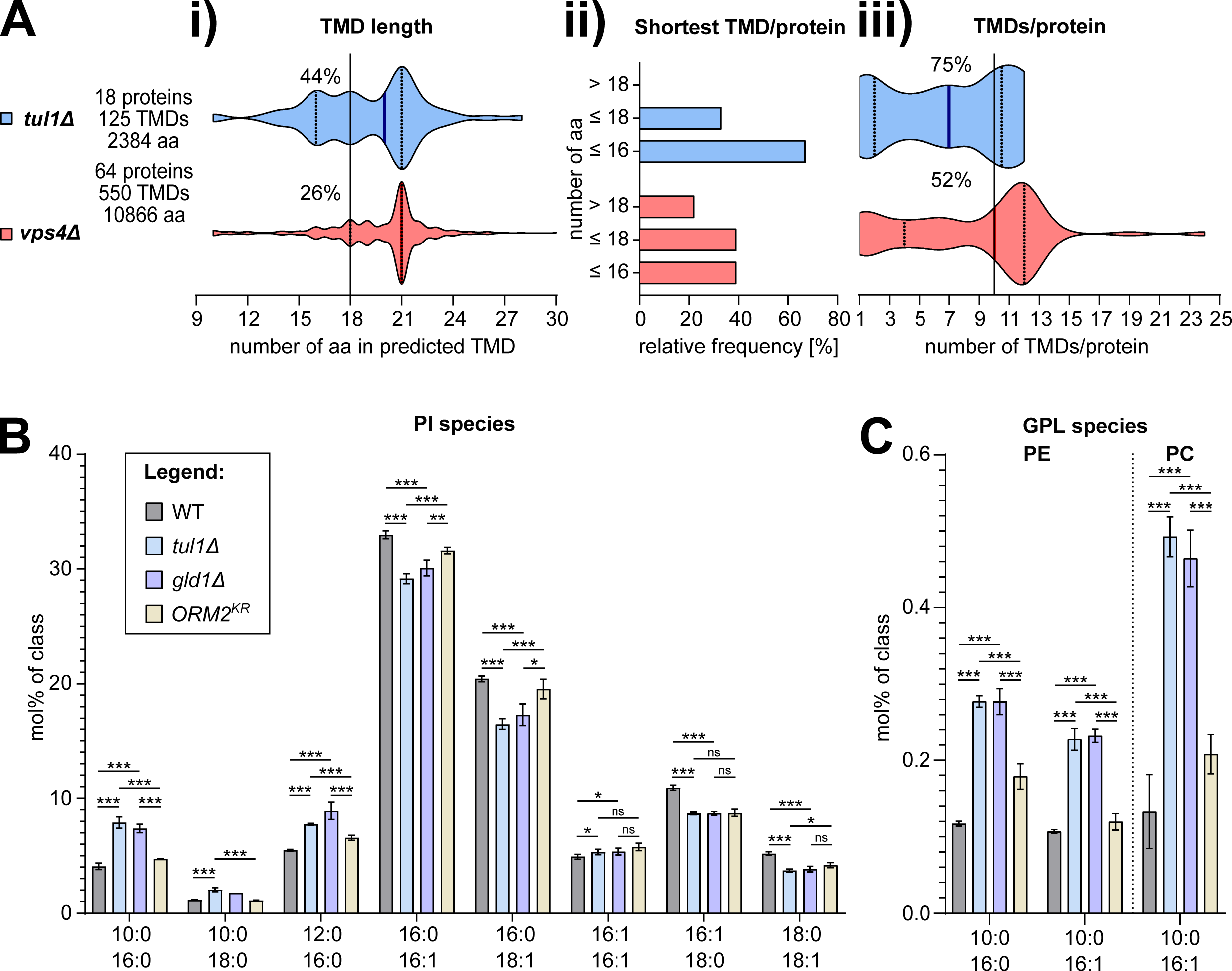
Transmembrane domains (TMDs) of ESCRT and EGAD mutants have different signatures. **(A)** Plots show the (i) number of amino acids in the predicted TMDs (**see Table S5**) in the upregulated proteins in *tul1Δ* and *vps4Δ*, (ii) the frequency of the shortest TMD/protein, (iii) the number of TMDs/protein mutant cells. Bold lines indicate median value, dotted lines indicate the first and third quartile. **(B, C)** Lipid abundances (measured using LC-MS) of **(B)** phosphatidylinositol (PI) species, **(C)** phosphatidylcholine (PC) and phosphatidylethanolamine (PE) species of lipid extracts from WT (dark gray), *tul1Δ* (light blue), *gld1Δ* (soft purple) and *ORM2^KR^* (pale yellow) mutant cells shown as mol% normalized to the respective lipid class (in mol%/class) shown as mean ± SD (n=4). P-values > 0.05 (ns), ≤ 0.05 (*), ≤ 0.01 (**) and ≤ 0.001 (***). See also **Figures S1, S2**.

It seemed that Dsc substrates were mainly ER and Golgi membrane proteins (Figure 1F), with relatively short TMDs (Figure 2A), while ESCRT clients were mainly PM proteins, with longer TMDs. These findings are consistent with the concept that TMDs have organelle-specific properties that match the lipid bilayer and sort proteins into ER/Golgi or PM lipid territories ^35-38^. Hence accumulating Dsc substrates might reside preferentially in ER/Golgi membranes with shorter and/or more unsaturated acyl chains glycerophospholipids (GPLs).

### Loss of the Gld1-Dsc complex caused specific membrane lipid adaptations

To analyze how the accumulation of Dsc substrates correlated with changes in membrane lipid composition, we used shotgun mass spectrometry-based lipidomics ^39^. We compared the lipid composition of WT cells to *tul1*Δ cells and *gld1Δ* cells. As an additional control we included cells in which the non-ubiquitinatable Orm2^KR^ mutant accumulated (but the Dsc complex remained functional) ^7^. Thereby, we identified changes that were caused specifically by the loss of the Golgi Gld1-Dsc complex, rather than by the Orm2 mediated inhibition of SPT activity or by the vacuolar Vld1-Dsc complex. Principal component analysis revealed that the lipid composition of *tul1Δ* cells was similar to *gld1Δ* cells, but differed from WT cells and from Orm2^KR^ cells (Figure S1A). We confirmed that in cells expressing Orm2^KR^, but also in *tul1Δ* cells, and in *gld1Δ* cells the levels of several ceramide, inositolphosphoryl ceramide, and mannosyl-inositolphosphoryl-ceramides (MIPC) were reduced (Figure S1B - E) ^7,28^. Hence these changes in sphingolipid metabolism were mainly caused by Orm2 accumulation and its inhibition of SPT activity.

The overall levels of the glycerophospholipids (GPL) classes were similar in WT, *tul1Δ* cells, *gld1Δ* cells, and in cells expressing Orm2^KR^ (Figure S2A). Yet, in *tul1Δ* and *gld1Δ* cells, there were specific changes in fatty acyl chain pairing, with an increase in shorter and asymmetric lipids. In *tul1Δ* and *gld1Δ* cells we measured an increase in PI, PE, PC, PS, and PA with shorter C-26 or C-28 fatty acyl chains (Figure 2B, C, S2B, C). The levels of C-26 almost doubled and C-28 increased by 50%. These GPL species paired fatty acyl chains that differed substantially in length: The PI-C-26 species showed a 2-fold increase in asymmetric acyl chains, and often both were saturated (PI 10:0 16:0) (Figure 2B). This effect was most pronounced for PI (Figure 2E), and was also observed for PE and PC (Figure 2C). Such asymmetric lipids had been reported earlier in budding yeast ^39^ and in fission yeast ^40^. Our results now showed, that the abundance of these asymmetric lipids increased with a loss of the Gld1-Dsc ubiquitin ligase complex and correlated with the accumulation of ER and Golgi proteins with short TMDs. Hence the Golgi Dsc complex maintained cellular protein and lipid composition.

### The heme oxygenase 1 (Hmx1) is ubiquitinated by the Dsc complex

To understand how the Dsc complex detected and degraded its substrates, we continued our analysis with the heme oxygenase 1 (Hmx1), the top-ranking Dsc substrate from our proteomic analysis (Figure 1B, D, E). Hmx1 is a conserved ER resident membrane protein ^41^ that is essential for the degradation of heme. The Alphafold (AF) model ^42,43^ of Hmx1 indicated a well-defined cytosolic enzymatic fold, and a single C-terminal tail-anchored TMD (Figure 3A).

**Figure 3:**
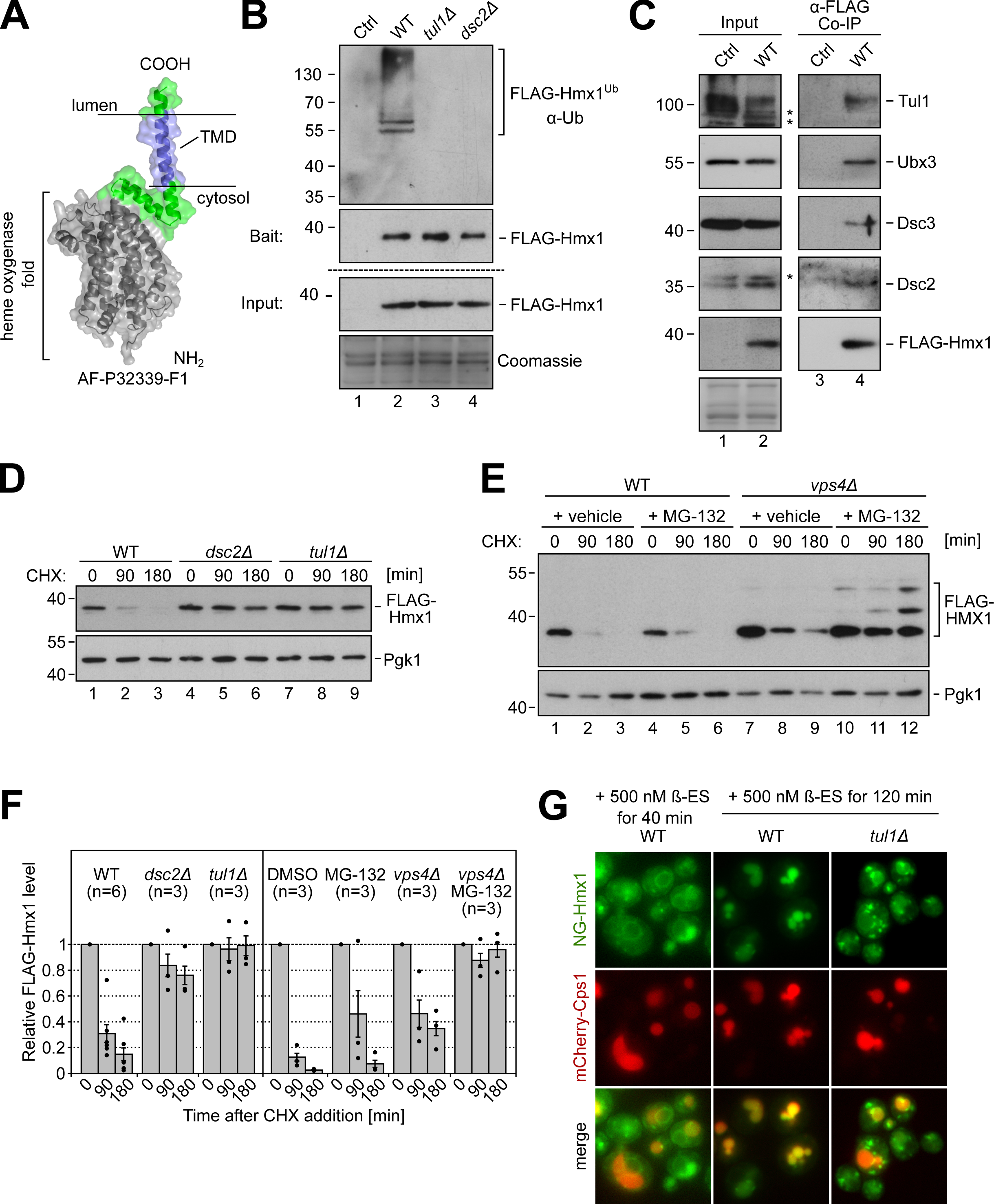
The heme oxygenase 1 (Hmx1) is ubiquitinated by the Dsc complex and degraded by ESCRT – and EGAD pathways. **(A)** AF model of Hmx1 (AF-P32339-F1). **(B-E)** SDS-PAGE and Western blot analysis with the indicated antibodies**: (B, C)** Input and elution of **(B)** denaturing or **(C)** non-denaturing FLAG-Hmx1 immunoprecipitations (IP) from indicated cells. Control (Ctrl) cells were untagged WT strains. ‘*’ unspecific antibody cross-reactions; **(D-E)** Total cells lysates of the indicated cells that were untreated (0 min) or treated with 50 µg/mL cycloheximide (CHX) to block protein synthesis for the indicated times. Pgk1 served as a loading control. **(F)** Densitometric quantification of FLAG-Hmx1 protein levels from Western blots of cell lysates of the indicated cells after the addition of CHX (n≥3), presented as mean ± standard deviation of the mean. **(G)** Live cell epifluorescence microscopy of the indicated cells, induced (500 nM ß-estradiol for the indicated times) mNeongreen-ALFA-Hmx1 (NG-Hmx1) (green) and mCherry-Cps1 (red). Scale bars 5µm. See also **Figure S3.**

To directly test whether Hmx1 was ubiquitinated in a Dsc complex dependent manner, we tagged endogenous Hmx1 with an N-terminal 3xFLAG tag (hereafter FLAG-Hmx1) and immunoprecipitated FLAG-Hmx1 from cell lysates of wild-type (WT) cells, *tul1*Δ, and *dsc2*Δ mutants under denaturing conditions. Ubiquitinated FLAG-Hmx1 was recovered from WT cells, but not from *tul1Δ* and *dsc2Δ* cells (Figure 3B). FLAG-Hmx1 migrated on an SDS–PAGE at around 38 kD. Ubiquitinated Hmx1 was detected at about 55–130 kD with laddering, suggesting polyubiquitination (Figure 3B). Under non-denaturing conditions, FLAG-Hmx1 co-immunoprecipitated the four subunits of the Dsc complex Tul1, Dsc2, Dsc3, and Ubx3 (Figure 3C). Hence, the Dsc complex interacted with Hmx1 and was essential for its polyubiquitination.

To determine how ubiquitinated FLAG-Hmx1 was degraded, we followed FLAG-Hmx1 protein turnover after blocking protein synthesis with cycloheximide (CHX). In WT cells, 70–80% of FLAG-Hmx1 was degraded after 90 min (Figure 3D, F). In cells lacking any component of the Golgi resident Dsc complex (*tul1*Δ, *dsc2*Δ, *ubx3*Δ, *dsc3*Δ or *gld1Δ*) the degradation of FLAG-Hmx1 was blocked (Figure 3D, S3A). Loss of the vacuolar Vld1-Dsc complex did not block Hmx1 turnover (Figure S3B). These results showed that the Gld1-Dsc complex ubiquitinated Hmx1 for degradation.

### Ubiquitinated Hmx1 can be degraded by ESCRT – and EGAD pathways

Our proteomics experiments (Figure 1) showed that Hmx1 accumulated in *tul1Δ* cells and in *vps4Δ* cells. Therefore, we next determined if Hmx1 was targeted into the EGAD pathway for proteasomal degradation or into the ESCRT pathway for vacuolar degradation.

Inhibition of the proteasome with MG-132 did not impair Hmx1 degradation (Figure 3E, F). Loss of ESCRT function (in *vps4*Δ cells) slowed down Hmx1 degradation, but also did not block it. Only when proteasomal activity was inhibited in *vps4Δ* mutants, was Hmx1 degradation blocked (Figure 3E, F). In these cells, we also observed the accumulation of higher molecular weight bands, possibly representing ubiquitinated Hmx1.

This demonstrated that the Dsc complex had the capacity to target Hmx1 either into EGAD- or ESCRT-pathways for degradation.

To follow the trafficking of Hmx1 out of the ER, we replaced endogenous Hmx1 with β-estradiol-inducible mNeonGreen-ALFA-Hmx1 (NG-Hmx1) ^44^. A signal for NG-Hmx1 was detected at the ER in WT cells, 40 min after the addition of β-estradiol (Figure 3G). After 120 min of induction, NG-Hmx1 was detected in the lumen of the vacuole in the majority of WT cells, where it co-localized with the ESCRT cargo mCherry-Cps1 (Figure 3G). During induction, the protein levels of full-length NG-Hmx1 reached a plateau at 90 min, and remained relatively constant (Figure S3C), while the protein levels of free NG increased, which was indicative of vacuolar proteolysis (Figure S3C). In *tul1Δ* cells, the majority of NG-Hmx1 accumulated in post-ER objects and on the limiting membrane of the vacuole after 120 min of induction (Figure 3G). Only a residual signal was detected in the lumen of the vacuole. Consistently, NG-Hmx1 protein levels increased more strongly in *tul1Δ* cells compared to WT cells and only little free NG was detected in *tul1Δ* mutants (Figure S3C). The small fraction of NG-Hmx1, that was sorted into vacuoles independently of the Dsc complex, might be the result of Rsp5 mediated ubiquitination of fluorescent proteins ^45^. Indeed, in *tul1Δ* cells, that additionally carried the hypomorphic *rsp5^G747E^* allele ^46^, NG-Hmx1 was no longer detected in vacuoles (Fig. S3D), whereas in the *rsp5^G747E^* cells, the Dsc complex targeted NG-Hmx1 into vacuoles.

These results suggested that Hmx1 was continuously exported from the ER to the Golgi, where the Dsc complex prevented its accumulation by ubiquitinating Hmx1 and targeting it preferentially into the ESCRT pathway for degradation inside vacuoles.

### Hmx1 is extracted from class E compartments for EGAD

It seemed that a large fraction of ubiquitinated Hmx1 was targeted into the ESCRT pathway for vacuolar degradation. Yet, loss of ESCRT function did not block Hmx1 degradation (Figure 3E, F, 4A). To dissect how the EGAD pathway contributed to Hmx1 degradation, we induced NG-Hmx1 expression for 90 min in *vps4*Δ cells and in *tul1*Δ *vps4*Δ double mutants. After the addition of CHX, the protein levels of mNG-Hmx1 substantially decreased in *vps4*Δ cells, but not in *tul1*Δ *vps4*Δ double mutants (Figure 4A). Of note, free NG (released by vacuolar degradation) was not detected, suggesting EGAD mediated proteasomal degradation. NG-Hmx1 accumulated on class E compartments in both *vps4*Δ cells and in *tul1*Δ *vps4*Δ double mutants, 90 min after the induction, together with the ESCRT client mCherry-Cps1 (Figure 4B). After blocking protein synthesis, the fluorescence signal for NG-Hmx1 markedly declined from class E compartments in *vps4*Δ cells and became difficult to detect, while the signal of mCherry-Cps1 remained unchanged (Figure 4B). In contrast, in *tul1*Δ *vps4*Δ double mutants the signal for NG-Hmx1 did not decrease on class E compartments, and still co-localized with mCherry-Cps1 on class E compartments (Figure 4B).

**Figure 4:**
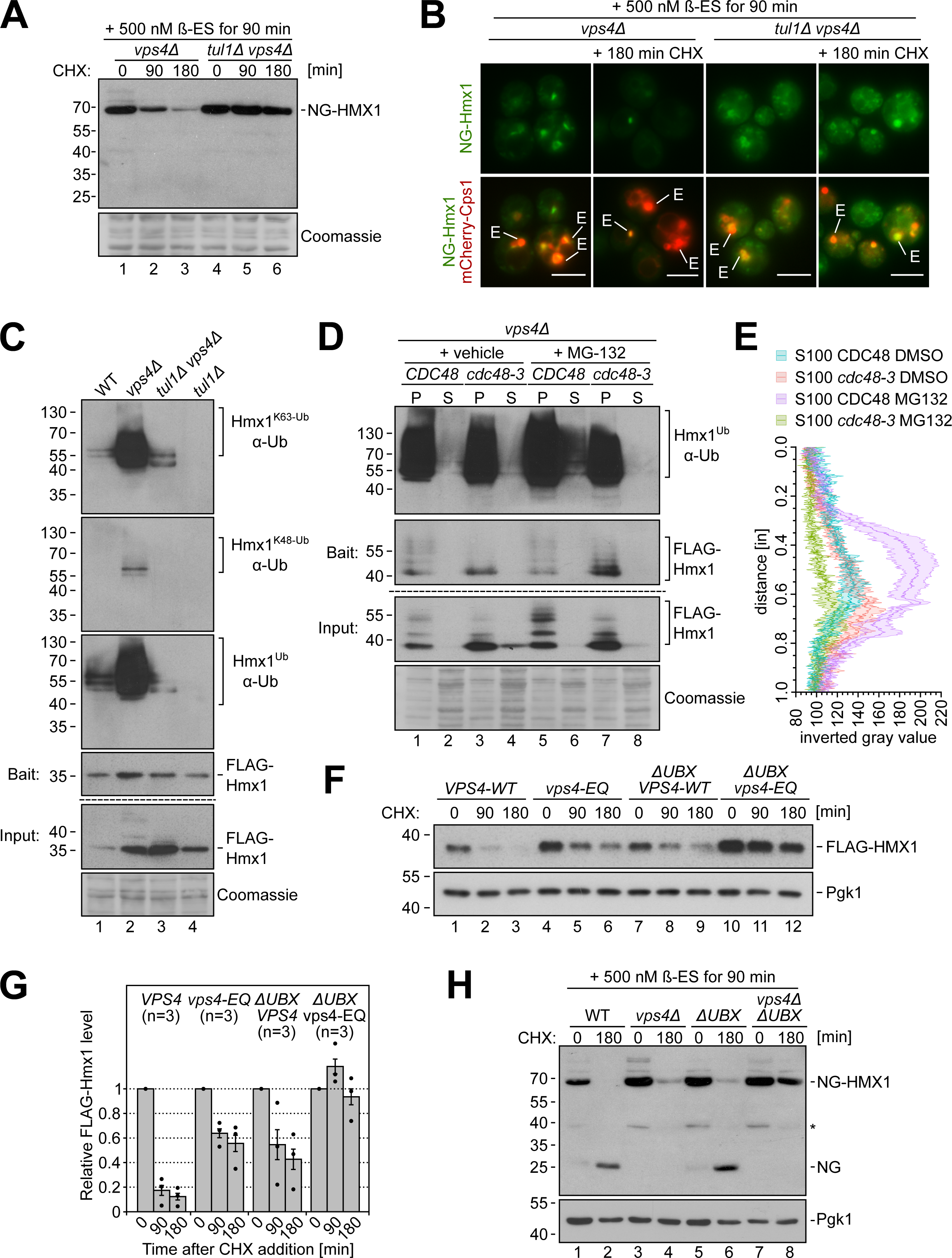
Hmx1 is extracted from class E compartments for EGAD. **(A-B)** NG-ALFA-Hmx1 (induced with 500 mM ß-estradiol (ß-ES) for 90 min) in *vps4Δ* and *tul1Δ vps4Δ* mutant cells. Cells were left untreated (0 min) or treated with CHX for the indicated time to block protein synthesis. **(A)** SDS-PAGE and Western blot analysis of total cell lysates with the indicated antibodies. Coomassie served as a loading control. **(B)** Live cells epifluorescence microscopy: NG-ALFA-Hmx1 (green) and Cherry-Cps1 (red). class E compartments are indicated; scale bars 5 µm. **(C-D)** SDS-PAGE and Western blot analysis with the indicated antibodies of input and elution of denaturing FLAG-Hmx1 immunoprecipitations (IP) from the indicated cells. **(C)** The two top panels: eluates from a 2^nd^ IP step with either K63 (top panel) - or K48-linkage specific nanobodies. Third panel: 1^st^ IP step with α-Flag antibody. **(D)** *CDC48* and *cdc48-3* mutants (in a *pdr5Δ vps4Δ* background) were left untreated or treated with 50 µM MG-132 to block proteasomal activity as indicated and subsequently subjected to subcellular fractionation. S100 and P100 fractions were subjected to denaturing FLAG-Hmx1 immunoprecipitations (IP). **(E)** Quantification of ubiquitinated FLAG-Hmx1 species from S100 fractions as plot profile, represented as mean ±95% confidence intervals of three individual line scans of S100 fractions. **(F,H)** SDS-PAGE and Western blot analysis of total cell extracts from the indicated cells. Cells were left untreated or treated with 50 µg/mL CHX for 90 min and 180 min. Pgk1 served as a loading control. For Orm2 see **Figure S4. (G)** Densitometric quantification of FLAG-Hmx1 protein levels 0, 90 and 180 min after the addition of CHX (n≥3), shown as mean ± standard deviation of the mean. **(H)** NG-Hmx1 expression was induced with 500 mM ß-estradiol (ß-ES) for 90 min prior to addition of 50 µg/mL cycloheximide (CHX). Pgk1 served as a loading control. ‘*’ indicates cross-reactive bands.

It seemed that the Dsc complex mediated the degradation of NG-Hmx1 from class E compartments via EGAD. This was consistent with our previous findings, that the Dsc complex localized to class E compartments for Orm2 degradation ^28^.

Since Cdc48 mainly extracts K48 polyubiquitinated proteins from membranes, we investigated polyubiquitin chain linkages on Hmx1. Therefore, we first immunoprecipitated Flag-Hmx1 from cells under denaturing conditions, followed by a second immunoprecipitation with nanobodies that are specific for either K63 or K48 polyubiquitin chains. In WT cells, Hmx1 was mainly K63 polyubiquitinated (Figure 4C). In *vps4Δ* cells, the levels of K63 polyubiquitination of Hmx1 increased, most likely because Hmx1 was no longer degraded along the ESCRT pathway. Now also K48 polyubiquitin chains on Hmx1 were detected. While the pool of K48 ubiquitinated Hmx1 was small, it was highly selective and mediated by the Dsc complex, since it was no longer detected in the *tul1*Δ *vps4*Δ cells. Also, K63 polyubiquitination was greatly reduced in the *tul1*Δ *vps4*Δ, but not completely abolished (Figure 4C). This residual polyubiquitination showed a different laddering. Yet, it was futile and Hmx1 was not degraded (Figure 4A, B).

Next, we tested if Cdc48 extracted ubiquitinated Hmx1 from class E compartments in *vps4*Δ cells. *vps4*Δ cells, left untreated or treated with proteasome inhibitor, were homogenized and the insoluble membrane fraction (P100) was separated from the soluble cytosolic fraction (S100) by centrifugation at 100,000 × *g* ^7,47,48^ (Figure 4D). FLAG-Hmx1 was immunoprecipitated from these fractions. Ubiquitinated Hmx1 predominantly accumulated in the membrane fraction (P100) (Figure 4D, E). As long as proteasomes were active or Cdc48 inactive, ubiquitinated Hmx1 was barely detected in the soluble fraction (Figure 4D, E). Yet, when proteasomal degradation was blocked and Cdc48 was active, a small fraction of membrane extracted polyubiquitinated Hmx1 was detected in the soluble fraction (S100) (Figure 4D, lane 6, E). We considered it likely that Cdc48 recruitment was facilitated by the UBX domain of Ubx3 ^18^. Indeed, in cells in which the UBX domain was deleted (*ubx3^ΔUBX^*), EGAD mediated Hmx1 degradation in ESCRT mutants was hampered (Figure 4F, lanes 10-12, G, H, lanes 7,8), while the ESCRT mediated degradation of Hmx1 was barley impaired (Figure 4F, lanes 7-9, G, H, lanes 5, 6). The appearance of a free NG band (at approximately 25kDa) in WT and *ubx3^ΔUBX^* cells suggested vacuolar turnover of NG-Hmx1 (Figure 4H, lane 2 & 6), and its absence indicated EGAD mediated proteasomal degradation in *vps4Δ* cells (Figure 4H, lane 4). In *ubx3^ΔUBX^*cells, also the degradation of Orm2 ^7^ was substantially delayed (Figure S4B, C) and GFP-Orm2 accumulated on post-ER compartments (Figure S4D).

Taken together these results showed that Dsc complex substrate Hmx1 was K63 polyubiquitinated for ESCRT mediated degradation in vacuoles, or K48 polyubiquitinated for EGAD mediated proteasomal degradation,

### Endogenous Yif1 is a Dsc complex substrate that is degraded by EGAD

To further explore how Dsc complex substrates are degraded, we next analyzed Yif1 turnover. Yif1 has been shown to be a Dsc complex substrate that is degraded by the ESCRT pathway ^24^. *YIF1* is an essential gene that encodes an evolutionarily conserved, integral membrane protein. Yif1 assembles with Yip1 and Yos1 into a complex that is required for the fusion of COPII vesicles with the Golgi ^49-51^. Yif1 was not quantified in our proteomics experiments.

To analyze whether Yif1 interacted with the Dsc complex, we replaced endogenous Yif1 with a C-terminally tagged Yif1-3xHA (Yif1-HA) (Figure 5A). Yif1-HA co-immunoprecipitated Tul1, Dsc2, Dsc3, and Ubx3 (Figure 5B). Next, we blocked protein translation to follow Yif1-HA turnover. In WT cells, 40% of Yif1-HA was degraded after 180 min and 60%-70% after 360 min (Figure 5C, S5C). In *tul1*Δ cells or when proteasomal degradation was blocked, the turnover of Yif1-HA was strongly impaired (Figure 5C, S5C). Furthermore, Yif1-HA was stabilized in *gld1*Δ mutants, while *vld1*Δ mutants still efficiently degraded Yif1-HA (Figure S5B, C). Next, we tested if the ubiquitin ligase activity of the Dsc complex was required for the degradation of Yif1-HA. Therefore, we mutated critical cysteines 748 and 751 in the RING domain of Tul1 to serine. The inactive mutant (Tul1^iRING^) impaired the degradation of Yif1-HA. In cells in which Cdc48 mediated membrane extraction (using the *cdc48-3* temperature sensitive allele or Ubx3^ΔUBX^ mutants) was hampered, Yif1-HA degradation was also impaired (Figure 5D, S5C). In contrast, in *pep4*Δ *prb1*Δ *prc1*Δ vacuolar protease triple mutant cells, as well as in *vps4Δ* cells, Yif1-HA degradation was not efficiently inhibited (Figure S5A, C). Likewise, when all ERAD E3 ubiquitin ligase complexes were disrupted (*doa10*Δ *hrd1*Δ *asi1*Δ), Yif1-HA was degraded as in WT cells (Figure S5A, C). When we replaced endogenous Yif1 with N-terminally eGFP tagged Yif1 (GFP-Yif1 expressed from an *ADH1* promoter), we detected GFP-Yif1 on discrete objects (most likely Golgi) in WT, which is consistent with its role in facilitating the fusion of COPII vesicles with the Golgi. GFP-Yif1 never co-localized Mup1-mCherry or mCherry-Cps1 in the lumen of the vacuole (Figure 5E, F). The localization of GFP-Yif1 was similar in *tul1Δ* cells. Dsc complex dependent vacuolar targeting of GFP-Yif1 ^24^ ^15^, was only observed when GFP-Yif1 was co-expressed in cells that still expressed endogenous Yif1 (Figure 5F).

**Figure 5:**
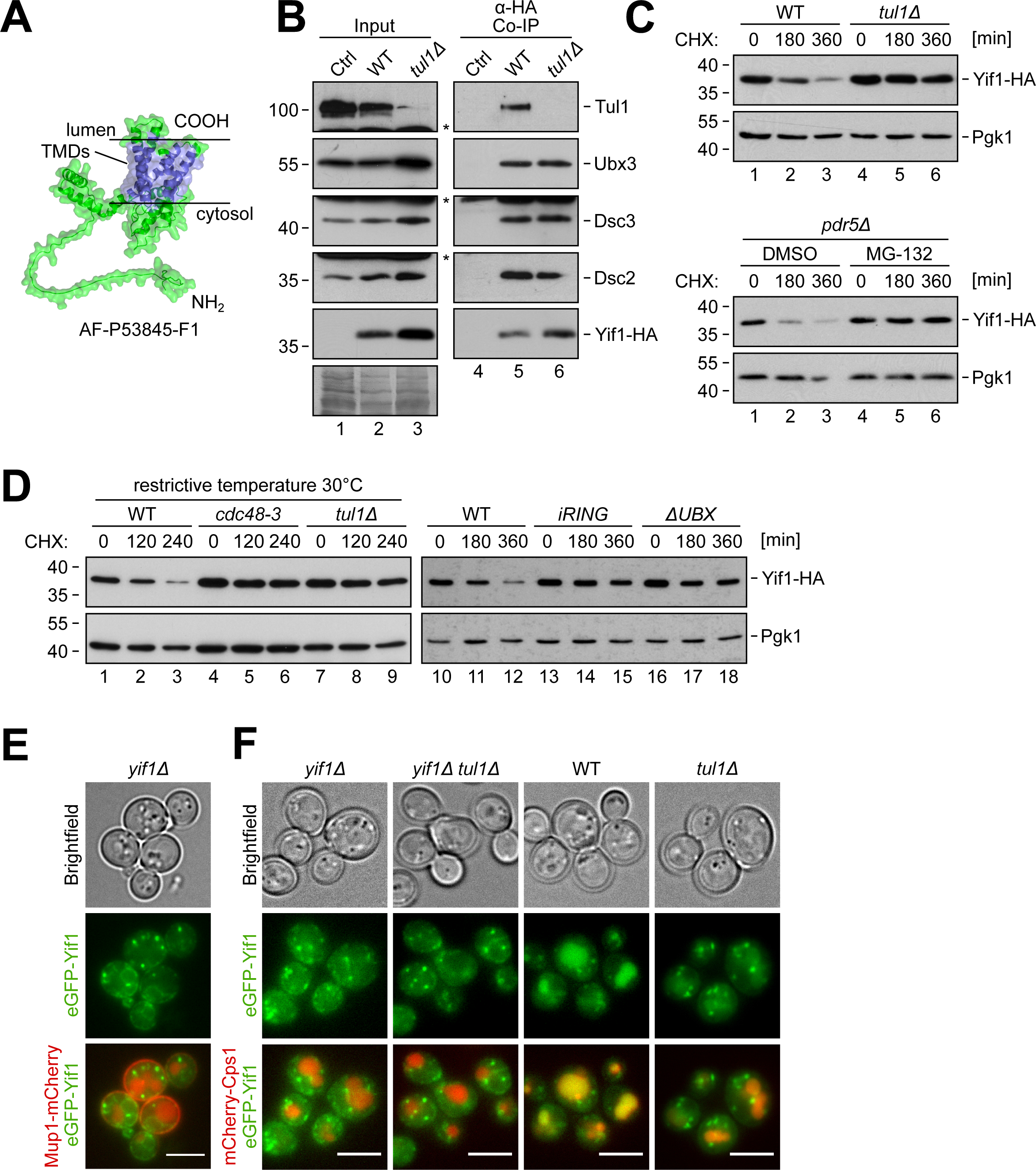
Endogenous Yif1 is a Dsc complex substrated and degraded by EGAD. **(A)** AF model of Yif1 (AF-P53845-F1). TMDs in blue. **(B-E)** SDS-PAGE and Western blot analysis with the indicated antibodies. **(B)** Input and elution from non-denaturing Yif1-HA co-immunoprecipitations from indicated cells. Control (Ctrl) cells were untagged WT strains. ‘*’ indicates cross-reactive bands; **(C-E)** Cells were left untreated (0 min) or treated with 50 µg/mL cycloheximide (CHX) for 180 min and 360 min. **(C)** Cells were incubated with 50 µM MG-132 or vehicle (DMSO) 10 min prior to the addition of CHX. **(D)** Cells were shifted to semi-permissive temperature (30°C). **(E, F)** Live cell epifluorescence microscopy of **(F)** *yif1Δ* cells expressing GFP-Yif1 (green) with Mup1-mCherry (red) and **(G)** of indicated cells expressing GFP-Yif1 (green) with mCherry-Cps1 (red). Scale bars 5 µm. See **Figure S5.**

These results were most consistent with a model in which a Gld1-Dsc complex mediated the degradation of endogenous Yif1 via EGAD, while excess Yif1 was targeted by the Dsc complex into the ESCRT pathway for vacuolar degradation.

### Dsc2 is required for substrate recognition

To characterize how the Dsc complex interacted with its substrates, we immunoprecipitated the substrates (FLAG-Orm2^KR^, FLAG-Hmx1 or Yif1-HA) from WT cells, or from *tul1*Δ, *dsc2*Δ, *dsc3*Δ, or *ubx3*Δ mutants (Figure 6A, Figure S6A). In WT cells, each substrate co-immunoprecipitated all subunits of the Dsc complex (Figure 6A, lanes 2, 8, 14). In contrast, in *dsc2*Δ cells, the interaction of the substrates with Tul1, Dsc3 or Ubx3 was strongly reduced (Figure 6A, lanes 4, 10, 17). In *ubx3*Δ cells, FLAG-Orm2, FLAG-Hmx1 and Yif1-HA co-immunoprecipitated reduced levels of Dsc2. FLAG-Hmx1 and Yif1-HA still interacted with Dsc3. In *dsc3*Δ cells, all substrates still co-immunoprecipitated Ubx3 and Dsc2, but the interaction with Tul1 was reduced. In *tul1Δ* cells, the interaction of the substrates with the remaining subunits of the Dsc complex was not affected.

**Figure 6:**
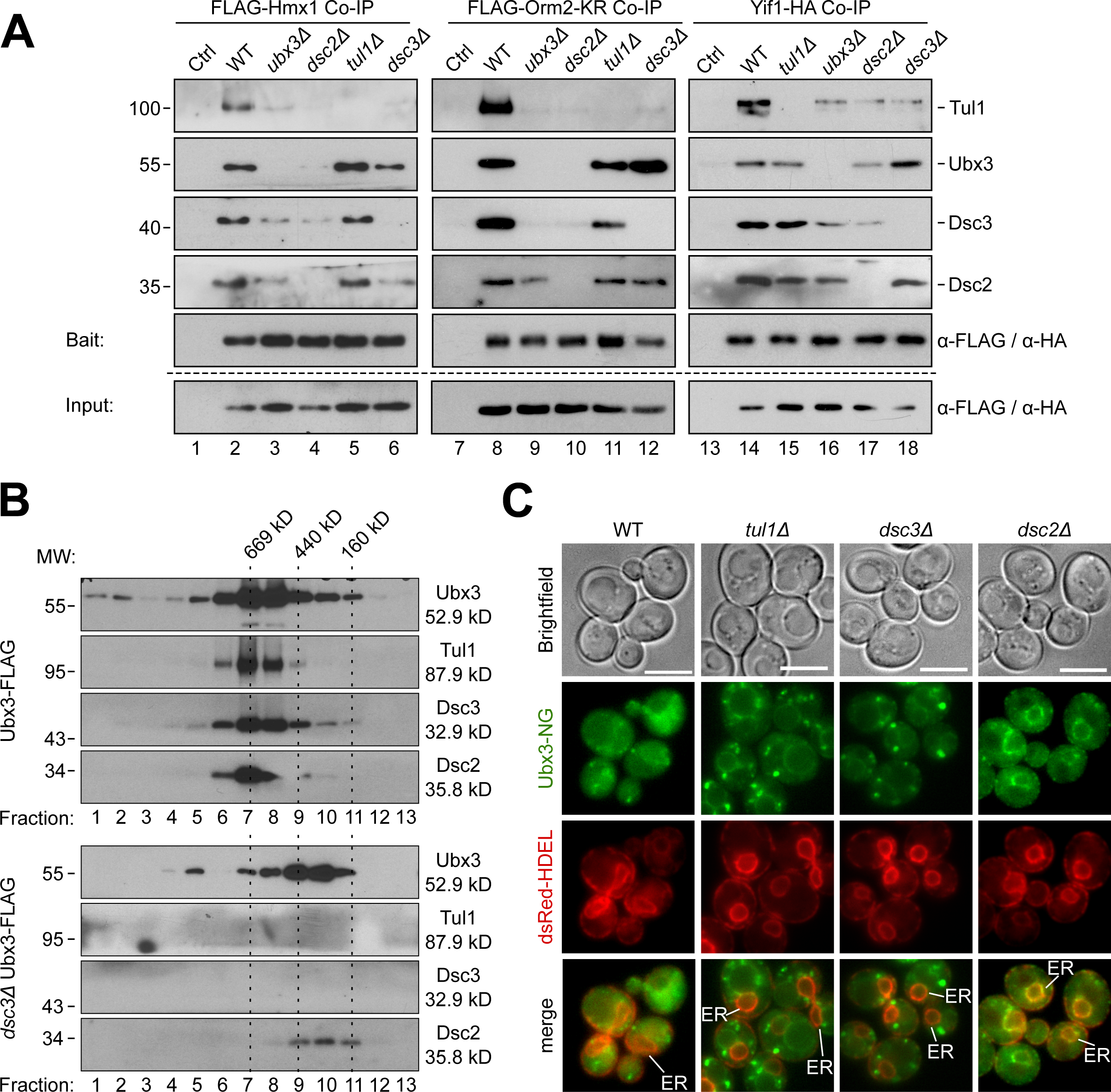
Dsc2 is required for substrate recognition. **(A-B)** SDS-PAGE and Western blot analysis with the indicated antibodies: **(A)** elution from non-denaturing FLAG-Hmx1, FLAG-Orm2-KR and Yif1-HA immunoprecipitations from the indicated cells. Control (Ctrl) cells were untagged WT strains. Inputs are shown in **Figure S6 (B)** Ubx3-Flag was immunoprecipitated from WT- or dsc3Δ cells and immunoprecipated proteins were subjected to size exclusion chromatography. Mr (in kDa) of the proteins and the standards are indicated. **(C)** Live cell epifluorescence of WT and the indicated mutants expressing Ubx3-mNG (green) with dsRED-HDEL (red). ER indicated, scale bars 5 µm.

These results suggested that Ubx3 and Dsc2 form a substrate recognition subcomplex. Consistently, when Ubx3-FLAG was immunoprecipitated from *dsc3Δ* cells, Tul1 interaction was no longer detected, but Dsc2 and Ubx3 formed a stable subcomplex. The immunoprecipitated Ubx3-Dsc2 subcomplex eluted from a size exclusion chromatography at a molecular weight that was smaller compared to that of the Dsc complex with all subunits (Figure 6B). Yet, the molecular weight of the Dsc complex and of the Ubx3-Dsc2 subcomplex were larger than expected for monomeric complexe. Yet, the interpretation is difficult due to glycosylation, micelle competition, shape of the Dsc complex, or unknown additional Dsc complex subunits. Along the same lines, live cell fluorescence microscopy revealed that Ubx3-NG was exported from the ER in *tul1*Δ and *dsc3*Δ cells, but remained trapped at the ER in *dsc2*Δ cells (Figure 6C). These results suggested that that Ubx3 and Dsc2 readily assembled at the ER for export.

Overall, these experiments indicated that Dsc3 recruited Ubx3 and Dsc2 to Tul1 which is consistent with earlier work ^14,15^. Moreover, Ubx3 and Dsc2, in particular Dsc2, were critical to interact with three different substrates.

### Dsc2 detects the TMD of Hmx1 as a degron

Dsc2 is a member of the rhomboid pseudoprotease family. These proteins can use their rhomboid like domain to locally perturb the membrane, which promotes membrane thinning for substrate engagement and/or retro-translocation ^52-55^. To analyze if Dsc2 would also have the potential to thin membranes, we used biomolecular dynamic modelling of the rhomboid like domain of Dsc2 (AF model: AF Q08232 F1). The Dsc2 rhomboid like domain was embedded into a lipid bilayer containing POPC (45%), POPE (45%) and cholesterol (10%) and molecular dynamics were simulated for 100ns. The simulation indicated that Dsc2 had the capacity to thin membranes between the L1 loop and TMD2 and TMD5 (Figure 7A, S7A, movie S1,2). Calculating the average center of mass between the membrane leaflets during the simulation and displaying it as a lipid density map revealed that, in this region of Dsc2, the lipid bilayer was thinned by about 5Å (Figure 7B, S7B, movie S3).

**Figure 7:**
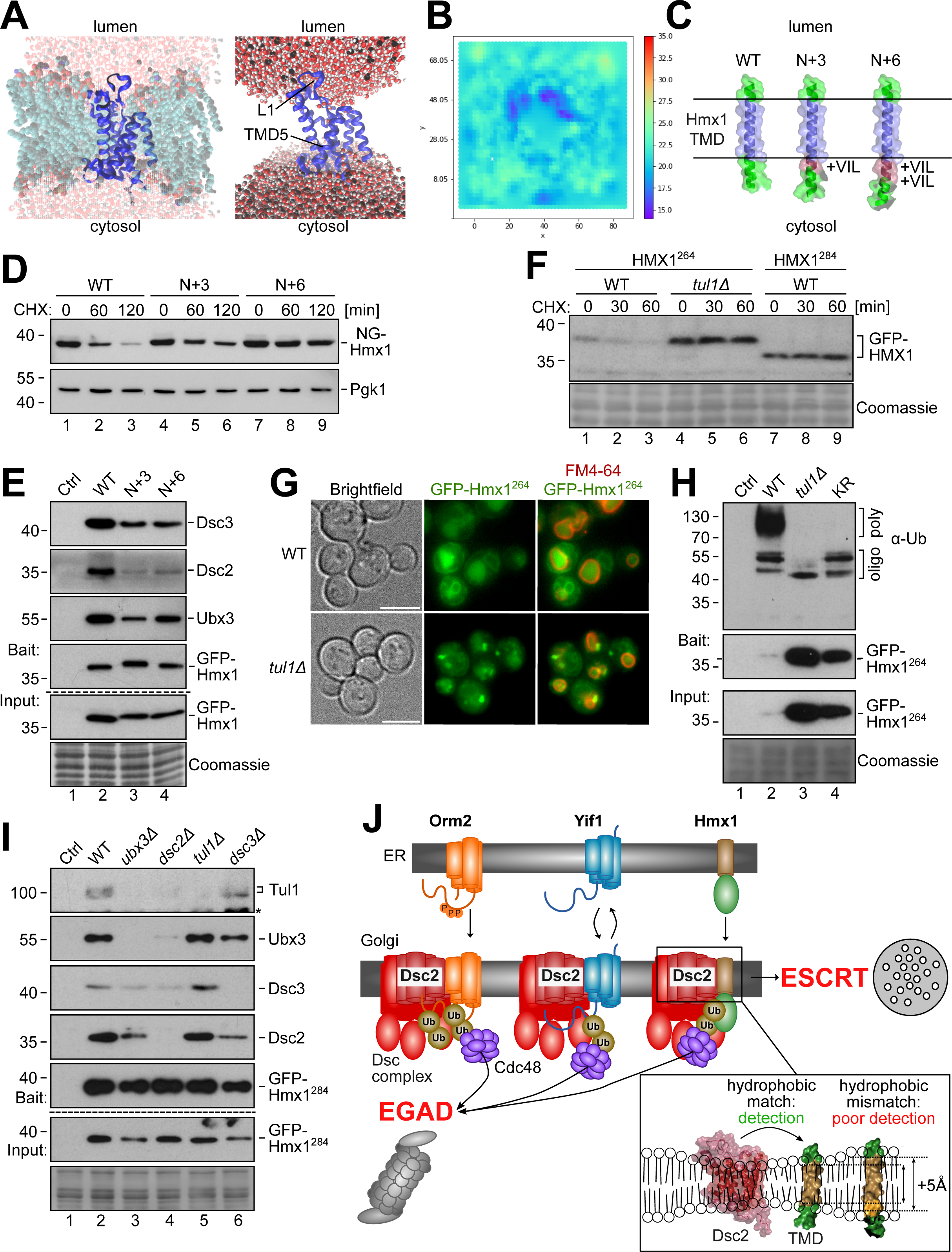
The TMD of Hmx1 is a degron for the Dsc complex. **(A)** Snapshot of the molecular dynamics simulation of the Dsc2 rhomboid domain showing the entire system (inculding lipids and water) or just the Dsc2 rhomboid domain with water in orthographic and perspective views. See also **movies S1, S2**. **(B)** Lipid density map displaying the average distances (in Å) of the center of mass in the lipid bilayer during the final converged 40ns of the simulation. See also **movie S3**. **(C)** AF models (AF-P32339-F1) showing the TMD of Hmx1 from amino acid (aa) 284-317 (WT), 284+3 aa (N+3), and 284+6 aa (N+6). Transmembrane domain (TMD) (blue) and extended α-helices (red) are colored as indicated. These constructs also encompass aa 264-317, but the cytosolic helix with aa 264-284 (including K265,269,282) is not shown. **(D-F, H, I)** SDS-PAGE and Western blot analysis with the indicated antibodies of the indicated mutants expressing the model substrates. Coomassie and Pgk1 served as a loading control. ‘*’ indicates cross-reactive bands. **(D, F)** Cells were left untreated (0 min) or treated with 50 µg/mL cycloheximide (CHX) to block protein synthesis for the indicated time points. **(D)** NG-Hmx1 expression was induced with 500 mM ß-estradiol (ß-ES) for 90 min prior to treatment with CHX. **(E)** Immunoprecipitations of different model substrates from *tul1Δ* cells. Control (Ctrl) cells were untagged WT cells. **(G)** Live cell epifluorescence of indicated cells expressing the model substrate GFP-Hmx1^264^ (green) with FM4-64 (red, vacuolar membrane). Scale bars 5 µm. **(H, I)** Input and elution of (H) denaturing GFP-Hmx1^264^ immunoprecipitations (IP) or (I) non-denaturing GFP-Hmx1^284^ from the indicated cells. Control (Ctrl) cells were untagged WT strains. See also **Figure S7. (J)** Summary of Dsc complex mediated degradation of orphaned transmembrane proteins.

To identify the degrons that could be detected by the Dsc2, we used Hmx1 as a model substrate because it has a single C-terminal TMD (18 aa, from aa residue 293 – 310). We generated a model substrate, Hmx1^264^ (Hmx1 amino acids residues 264 – 317), which included the C-terminal TMD (Figure 7C) and a short cytosolic α-helix with three lysine residues (K265, K269, K282) (not shown). Because TMD length was a discriminator for Dsc complex substrates and because Dsc2 has the potential to thin membranes by about 5 Å, we generated two additional model substrates in which we extended N-terminally the hydrophobic length of the TMD by addition of 3 (LIV) or 6 amino acids (LIVLIV) (Figure 7C), which adds an additional pitch of around 4,5 Å and 9 Å (Figure 7C). Hmx1^264^ (WT), Hmx1^264+3^ (N+3) or Hmx1^264+6^ (N+6) (Figure 7C), were N-terminally fused to mNeonGreen-ALFA-(NG)-tags or GFP-ALFA-(GFP)-tags. When the model substrates where constitutively expressed (from a strong TDH3 promotor), the protein levels of GPF-Hmx1^264^ were low (Figure S7C), while the protein levels of the longer model substrates were higher (Figure S7C). The clipped GFP suggested that these proteins were still sorted into vacuoles, but not as efficiently as the WT model substrate. To directly compare their turnover, we induced the expression of the model substrates for 90 minutes with β−estradiol. After induction, the protein levels NG-Hmx1^264^, and of the longer model substrates N+3 and N+6 were similar, but NG-Hmx1^264^ protein levels rapidly declined within 60-120 minutes (Figure 7D, lanes 1-3), while the degradation of the longer model substrates N+3 and N+6 was markedly delayed. To compare the interaction of these model substrates with the Dsc complex, we immunoprecipitated GPF-Hmx1^264^, N+3 and N+6 from *tul1Δ* cells. In *tul1Δ* cells, the steady state protein levels of the different model substrates were comparable (Figure 7E). Notably, the longer model substrates interacted less efficiently with Dsc2, Dsc3 and Ubx3 (Figure 7E).

To pinpoint if the TMD of Hmx1 contained a Dsc complex specific degron, we determined the turnover of GPF-Hmx1^264^ in WT cells and in *tul1*Δ cells. In WT cells the protein levels of GFP-Hmx1^264^ were low and declined within 60 min (Figure 7F). In contrast, in *tul1*Δ cells, GFP-Hmx1^264^ levels were readily elevated and the protein was not degraded (Figure 7F). Consistently, live cell fluorescence microscopy showed, that GFP-Hmx1^264^ or pHluorin-Hmx1^264^ accumulated in dot-like objects, probably Golgi and endosomes, and on the limiting membrane of vacuoles in *tul1Δ* cells (Figure 7G, S7D). In WT cells, GFP-Hmx1^264^ localized to the ER and to the lumen of FM4-64 labeled vacuoles (Figure 7G) and pHluorin-Hmx1^264^ was quenched inside vacuoles but detected at nuclear and cortical ER (Figure S7D).

The cytosolic K265, K269, K282 of GFP-Hmx1^264^ were heavily polyubiquitinated in a Dsc complex dependent manner. When the three cytosolic lysine residues were mutated to arginine (K265,269,282R), GFP-Hmx^264-KR^ accumulated, but polyubiquitination was no longer detected (Figure 7H), similar to *tul1*Δ cells. Only a small fraction of apparently mono or oligo-ubiquitinated GFP-Hmx1^264^ was detected (Figure 7H). Thus, K265, K269 and K282 are the major targets for Dsc complex mediated polyubiquitination and degradation.

GFP-Hmx1^264^ was heavily K48 polyubiquitinated in WT cells and in *vps4*Δ cells by Tul1 (Figure S7E, compare lanes 1 and 2 with lane 3). This Tul1 mediated K48 polyubiquitination of GFP-Hmx1^264^, was essential for EGAD mediated degradation from class E compartments in ESCRT mutants (Figure S7F, G). In *vps4Δ* cells, GFP-Hmx1^264^ was efficiently degraded (Figure S7F), while in *vps4*Δ *tul1*Δ cells, GFP-Hmx1^264^ was no longer degraded and, thus, accumulated (Figure S7F). Consistently, in cells expressing Vps4^E233Q^, GFP-Hmx1^264^ was mainly detected at the ER, with little or no co-localization with FM4-64 on class E compartments (Figure S7G). Yet, when the UBX domain of Ubx3 was deleted, GFP-Hmx1^264^ could no longer be efficiently extracted from class E compartments, and accumulated with FM4-64 on class E compartments (Figure S7G).

Finally, we analyzed the shorter model substrate GFP-Hmx1^284^, which consistent only of the C-terminal TMD (18 aa, from aa residue 293 – 310), but lacked the short cytosolic α-helix with three lysine residues (K265, K269, K282). The protein levels of the lysine-less model substrate GFP-Hmx1^284^ were high in WT cells and did not decrease (Figure 7F, lanes 7-9). Yet, this short model substrate GFP-Hmx1^284^ was sufficient to interact with the Dsc complex. Non-denaturing immunoprecipitation experiments showed that GFP-Hmx1^284^ efficiently co-immunoprecipitated all subunits of the Dsc complex (Figure 7I, lanes 2). This interaction required Dsc2, but not Tul1 (Figure 7I, lanes 4 and 5).

Our results demonstrated that the TMD of Hmx1 is a degron for the Dsc complex and its hydrophobic length was assessed by Dsc2, for determined efficient substrate engagement. Thereby, the Dsc complex detects orphaned membrane proteins (Figure 7J), prevents their accumulation at the Golgi and limits their spreading to other organelles.

## Discussion

Our work shows that the Dsc ubiquitin ligase complex plays a central role in membrane quality control at the Golgi and endosomes, where it detects orphaned proteins and ubiquitinates them for degradation (Figure 7J): This model includes the following steps: (1) The Ubx3-Dsc2 assemble at the ER and are required for substrate recognition at the Golgi. (2) There, Dsc2 can locally thin the membrane and is essential to interact with proteins that expose short TMDs as degrons. (3) Substrates with such degrons are typically ER and Golgi membrane proteins. (4) Dsc3 connects Ubx3-Dsc2 with the E3 ubiquitin ligase Tul1. (5) Tul1 coordinates the E2 enzyme Ubc4 to conjugate ubiquitin onto the substrates.

The ubiquitination of all known Dsc substrates (including PE) appears to require the canonical enzymes Uba1 (E1), Ubc4/5 (E2), and Tul1 (E3) ^13,16,24,27^. Since Ubc4 is required for ESCRT and EGAD mediated degradation of Dsc substrates, we propose that Tul1-Ubc4 sets a priming mono- or oligo-ubiquitination, that can be extended into different multi-or polyubiquitin chains. For EGAD, the conjugated ubiquitin residues are elongated into K48 polyubiquitin chains. Cdc48 could bring along Ufd2 as an E4 ubiquitin ligase to extend the priming ubiquitination mediated by Ubc4-Tul1 into a K48 polyubiquitin chain. This would reinforce Cdc48 mediated membrane extraction or subsequent proteasomal degradation or both ^56-60^. Orm2 and endogenous Yif1 follow this pathway.

For ESCRT mediated vacuolar degradation, ubiquitinated residues are either not elongated or elongated into K63 polyubiquitin chains. Elongation could involve other E3 ubiquitin ligases such as Rsp5 or Pib1 ^61,62^. This would support the selectivity of the ESCRT-0 complex towards K63 polyubiquitin chains ^63,64^. Thus, the Dsc complex, together with other ubiquitin ligases, may work sequentially in Golgi PQC, to cover different degrons and ‘ubiquitination zones’ ^61,65^.

The formation of the different polyubiquitin chains might depend on how the Dsc complex interacts with its substrates. Substrates with shorter TMDs may be better hydrophobically matched with Dsc2, which could increase the dwell time and, thus, successful Cdc48 engagement and membrane extraction, while faster dissociation from Dsc2 could result in the sorting of the ubiquitinated substrates into the ESCRT pathway.

The three Dsc complex substrates, Orm2, Yif1, and Hmx1, are ER and Golgi proteins with relatively short TMDs and minimal luminal or cytosolic loops. Their TMDs harbor degron features that are recognized by Dsc2, most likely at the trans-Golgi. Degron detection might be facilitated by the capacity of Dsc2 to locally perturb the membrane ^52,53^, similar to the function of Derlins in ERAD ^54,55^. Dsc2 mediated thinning of the lipid bilayer could promote hydrophobic matching with the shorter TMD degrons and/or additionally destabilize TMDs to overcome the energetic barrier during retro-translocation and Cdc48 mediated membrane extraction ^66-68^.

For Hmx1, the tail-anchored TMD functions as a degron that is detected by Dsc2 at the Golgi. How could Dsc2 detect orphaned Orm2? At the ER, Orm2 interacts with the membrane-spanning helix of Lcb1 of the SPT via its short TMD3 (14 amino acids) and TMD4 (18 amino acids) ^69^ ^70^. Once Orm2 is exported from the ER into the Golgi, it no longer interacts with Lcb1, and the TMD3 and TMD4 would be exposed and detected by Dsc2 for EGAD. A similar case could be made for Yif1 ^49-51^. Based on AF models of the Yif1-Yip1 complex in ModelArchive ^71^, the Yip1-Yif1 interaction occurs along TMD1 of Yif1 which is predicted to encompass 18 amino acids. For orphaned Yif1, the TMD1 might be a degron that is detected by Dsc2. Alternatively, exposure of the degron might occur when Yif1 is idle or when Yif1 departs from its regular ER-*cis-*Golgi route into the *trans-*Golgi.

The spatial separation of the Dsc complex to the Golgi prevents the premature degradation of free Orm2 and Yif1 at the ER. This could improve the efficiency of Orm2-SPT or Yif1-Yip1 complex assembly at the ER and, at the same time, reduce the level of orphaned Orm2 and Yif1 at the Golgi. A similar concept was proposed for the quality control of unassembled membrane proteins by Asi-ERAD at the inner nuclear membrane ^72-74^.

It remains to be tested if the degradation of human orthologues of Orm2 (ORMDL1-3) ^75^, Yif1 (YIPF3), and Hmx1 (HO-1) ^76,77^ is conceptually similar. However, it is clear that ORMDL and HO-1 are ubiquitinated, membrane extracted and degraded by proteasomes. The degradation of HO-1, requires MARCH6/TEB4 and TRC8 in a redundant manner together with the signal peptide peptidase (SSP) ^76^. The same study also found ORMDL2 among the upregulated proteins in MARCH6/TRC8 double knockout cells.

In Dsc complex deficient cells, Orm2, Yif1 and Hmx1 accumulate at Golgi, but also on endosomes and the limiting membrane of vacuoles. Interestingly, they were always excluded from the plasma membrane (PM) and also from other membrane regions of increasing thickness, presumably by hydrophobic mismatching ^37,38,78-82^. Hence, our results are in agreement with previous work that proposed that physical properties of TMDs, such as length, hydrophobicity, and volume, function as sorting signals within the lipid bilayers of eukaryotic cells and must be matched by the hydrophobic core of the lipid bilayer ^35,40,80,83-88^. Our analysis suggests that, these TMD properties also function as degrons to flag orphaned proteins. Almost reciprocally, the accumulation of Dsc substrates causes an increase in shorter, saturated and asymmetric GPL. Such saturated asymmetric lipid species have been detected recently in fission and budding yeast ^39,40^. *S. japonicus*, but not *S. pombe,* shows an enrichment of proteins with relatively short TMDs alongside asymmetric saturated GPL ^40^. Membranes with such GPLs could accommodate the accumulating Dsc substrates by improving hydrophobic matching while reducing membrane permeability ^89,90^.

Thus, the Dsc complex may be ideally localized at the Golgi for membrane quality control. There, it integrates lipid bilayer thickness with TMD length. Such hydrophobic TMD matching could provide a general means of PQC machineries to detect and degrade orphaned protein, and limit their spreading across the cell.

## Limitations of this study

While our work indicates that the Dsc complex detects TMD degrons by hydrophobic matching, we do not yet understand how the downstream EGAD or ESCRT pathways are selected for degradation. Moreover, to better understand the molecular mechanism of hydrophobic matching, the process of Dsc2 mediated substrate recognition must be reconstituted in an *in vitro* system.

## Acknowledgments

We thank Snezhana Oliferenko, Hesso Farhan, Chris Dunworth and Lukas A Huber for critically reading the manuscript, Ming Li, Peter Espenshade and Scott Emr for reagents. This research was funded in part by the Austrian Science Fund (FWF) (10.55776/P32161, 10.55776/P34907, 10.55776/DOC82 to DT, and 10.55776/P36187 to OS), by a Lipotype lipidomics excellence award (LEA 2019) to OS, by a Luxembourg National Research Fund (FNR): Grant #13571826 to YW, and by European Union’s Horizon 2020 research and innovation program under the Marie Skłodowska-Curie grant agreement No. 847681 (to KRL). For open access purposes, the author has applied a CC BY public copyright license to any author accepted manuscript version arising from this submission.

## Author contributions

YW, SIS, designed and performed the vast majority of experiments and analyzed data, with support from MW, VN, MT, SW. BS and LK conducted quantitative proteomics, OS prepared samples for lipidomics analysis. XT and KR performed and analyzed MDS. DT and YW wrote the manuscript. DT conceived and supervised the study.

---

**Figure S1: Impact on lipid and sphingolipid synthesis in *tul1Δ*, *gld1Δ* and *ORM2^KR^* mutant cells.**

**(A)** Principal component analysis of lipid extracts from WT (dark gray), *tul1Δ* (light blue), *gld1Δ* (soft purple) and *ORM2^KR^* (pale yellow) mutant cells. **(B)** Abundance of ceramides (Cer), inositolphosphoryl-ceramides (IPC), mannosyl-inositolphosphoryl-ceramides (MIPC), mannosyl-diinositolphosphoceramides (M(IP)2C) classes of lipid extracts from WT (dark gray), *tul1Δ* (light blue), *gld1Δ* (soft purple) and *ORM2^KR^* (pale yellow) mutant cells normalized to the total lipid content. **(C-E)** Abundance of different **(C)** ceramide, **(D)** IPC and **(E)** MIPC species of lipid extracts from WT (dark gray), *tul1Δ* (light blue), *gld1Δ* (soft purple) and *ORM2^KR^*(pale yellow) mutant cells shown as mol% normalized to the respective lipid class. Lipid extracts were measured using LC-MS. Data is presented as mean ± standard deviation from four independent experiments (n=4) and legend for all graphs is shown on the right-hand side of the figure. Downregulated species are labeled in red. P-values are flagged with > 0.05 (ns), ≤ 0.05 (*), ≤ 0.01 (**) and ≤ 0.001 (***).

**Figure S2: Impact on glycerophospholipids (GPLs) acyl chain length and species in *tul1Δ*, *gld1Δ* and *ORM2^KR^* mutant cells.**

**(A)** Abundance (in % of total lipids) of phosphatidic acid (PA), phosphatidylcholine (PC), phosphatidylethanolamine (PE), phosphatidylinositol (PI) and phosphatidylserine (PS) of lipid extracts from WT (dark gray), *tul1Δ* (light blue), *gld1Δ* (soft purple) and *ORM2^KR^* (pale yellow) mutant cells normalized to the total lipid content. **(B-C)** Abundances (in mol%/class) of different **(B)** PI acyl chain length, **(C)** PS, PA, PC and PE acyl chain length.

**Figure S3: Hmx1 is a substrate of the Gld1-Dsc complex.**

**(A-C)** SDS-PAGE and Western blot analysis with the indicated antibodies of WT cells and the indicated mutants that were either untreated (0 min) or treated with 50 µg/mL cycloheximide (CHX) to block protein synthesis for the indicated time points. Coomassie and Pgk1 served as a loading control. **(C)** NG-Hmx1 expression was induced with 500 nM β-ES and samples were collected at the indicated time points. **(D)** Densitometric quantification of NG-Hmx1 and free NG protein levels. Each experiment was repeated three times, the sum of NG-Hmx1 and free NG levels were set to 1, and their respective fraction was related to the sum. Data are presented as mean ± standard deviation of the mean (SEM). **(E)** Epifluorescence microscopy of living *rsp5^G747E^* and *tul1Δ rsp5^G747E^* mutant cells expressing inducible NG-Hmx1 (green). NG-Hmx1 expression was induced with 500 nM β-ES and imaged after 300 min. Scale bars 5 µm.

**Figure S4: The UBX domain of Ubx3 is required for Orm2 degradation**

**(A)** SDS-PAGE and Western blot analysis with the indicated antibodies of WT cells and the indicated mutants that were either untreated (0 min) or treated with 50 µg/mL cycloheximide (CHX) to block protein synthesis for the indicated time points. FLAG-Orm2 was co-expressed from an episomal plasmid. **(B)** Densitometric quantification of FLAG-Orm2 protein levels from Western blots of cell lysates. Each experiment was repeated at least three times, protein levels were normalized to Pgk1 loading controls, and each time-point was related to t=0 min (set to 1). Data are presented as mean ± standard deviation of the mean (SEM). **(C)** Epifluorescence microscopy of living WT and *ubx3^ΔUBX^* cells co-expressing GFP-Orm2 (green) from an episomal plasmid. Scale bars 5 µm.

**Figure S5: Yif1-HA is a Gld1-Dsc substrate**

**(A-B)** SDS-PAGE and Western blot analysis with the indicated antibodies of WT cells and the indicated mutants that were either untreated (0 min) or treated with 50 µg/mL CHX for the indicated time points to block protein synthesis for the indicated time points. **(C)** Densitometric quantification of Yif1-HA protein levels from Western blots of cell lysates from (Figure 5 C, D and Figure S5A). Each experiment was repeated at least three times, protein levels were normalized to Pgk1 loading controls, and each time-point was related to t=0 min (set to 1).

**Figure S6: Inputs of substrate co-immunoprecipitations**

**(A)** SDS-PAGE and Western blot analysis with the indicated antibodies of inputs from non-denaturing FLAG-Hmx1, non-ubiquitinatable FLAG-Orm2-KR and Yif1-HA co-immunoprecipitations from (Figure 6A) from WT cells and the indicated mutants. Control (Ctrl) cells were untagged WT strains. Coomassie served as loading control. ‘*’ indicates the correct band.

**Figure S7: The TMD of Hmx1 is a Dsc complex degron**

**(A)** Additional snapshots of the molecular dynamics simulation of the Dsc2 rhomboid domain showing the entire system (Dsc2 rhomboid domain, lipids and water) or just with the Dsc2 rhomboid domain and water in perspective and orthographic views. **(B)** Overlay of the lipid density map displaying the distance of the average center of mass in the lipid bilayer over the final 40ns of the simulation with a top view of the Dsc2 rhomboid domain (from the luminal site) and the lipids. **(C)** SDS-PAGE and Western blot analysis with the indicated antibodies of whole cell lysates from WT cells expressing the indicated versions of the Hmx1 TMD. **(D)** Epifluorescence and phase contrast microscopy of living WT and *tul1Δ* mutants expressing truncated pHluorine-Hmx1 from residue 264-317 (green) with mCherry-Cps1 (red, MVB cargo). Scale bars 5 µm. **(E)** SDS-PAGE and Western blot analysis with the indicated antibodies of input and elution of denaturing eGFP-ALFA-Hmx1^264^ immunoprecipitations (IP) from WT cells and the indicated mutants. Coomassie served as a loading control. The two top panels show the eluates from a 2^nd^ IP step with K48- or K63-linkage specific nanobodies decorated with an anti-ubiquitin antibody. **(F)** Epifluorescence and phase contrast microscopy of living WT and *ubx3^ΔUBX^ (ΔUBX)* mutants expressing truncated GFP-Hmx1 from residue 264-317. Cells additionally expressed either *VPS4* or *vps4-E233Q* (to block vacuolar sorting) from an episomal plasmid. Panel shows GFP-Hmx1^264^ (green) with FM4-64 (red, vacuolar membrane). E indicates class E compartments; scale bars 5 µm. **(G)** SDS-PAGE and Western blot analysis with the indicated antibodies of WT cells and *tul1Δ* expressing truncated eGFP-ALFA-Hmx1 from residue 264-317. Cells were left untreated (0 min) or treated with 50 µg/mL cycloheximide (CHX) to block protein synthesis for 45 min and 90 min. Coomassie served as a loading control.

**Table S6.**
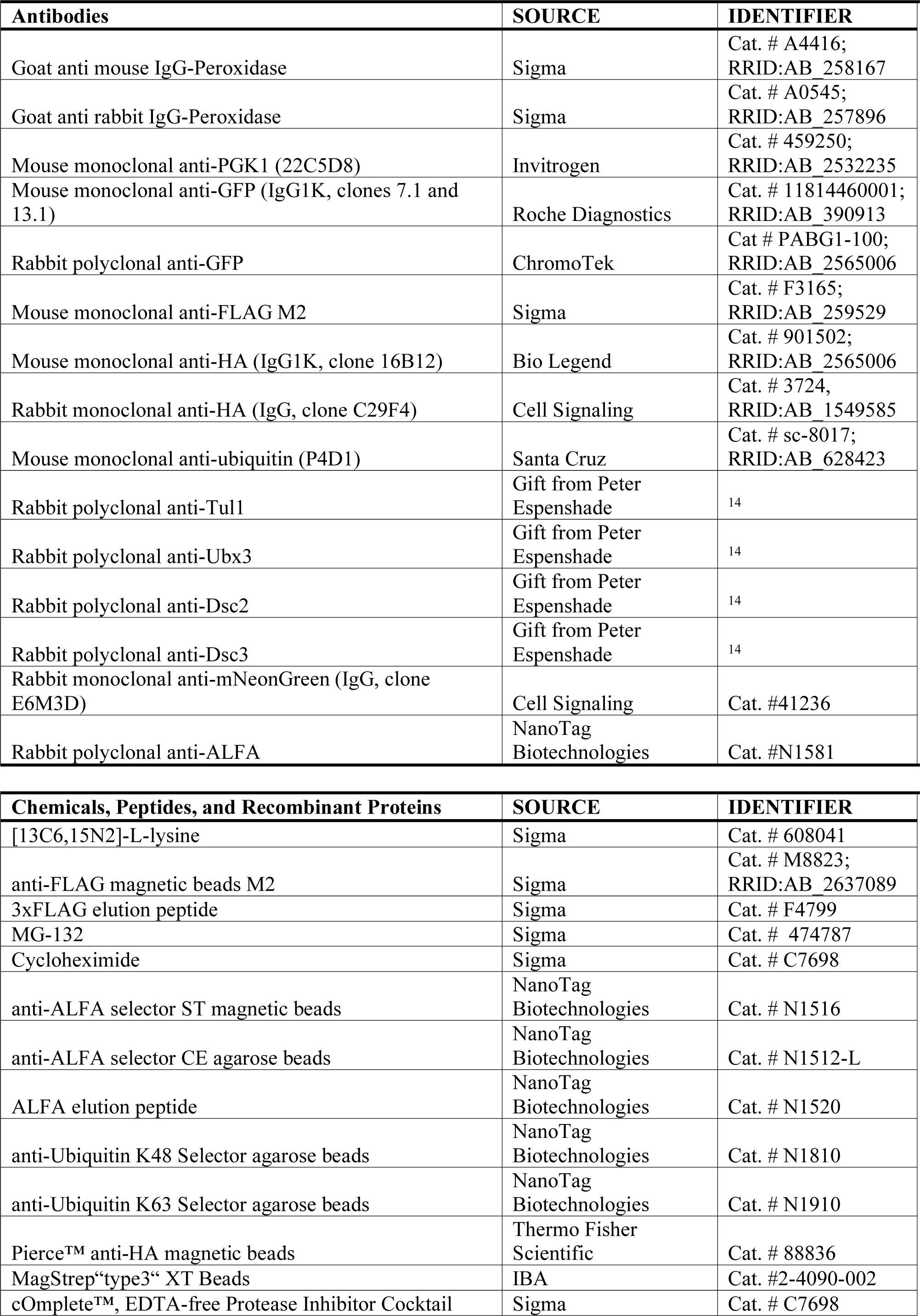
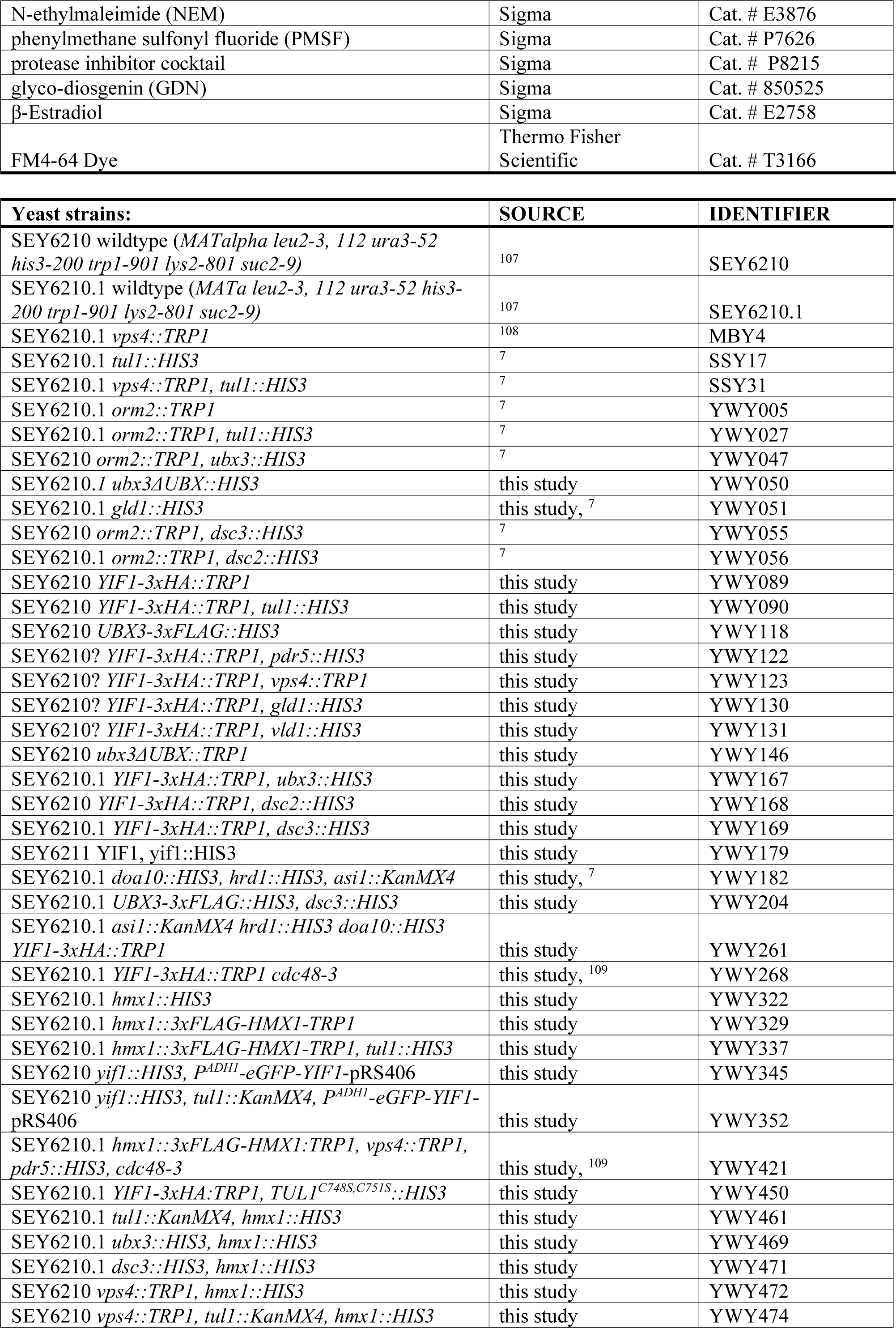
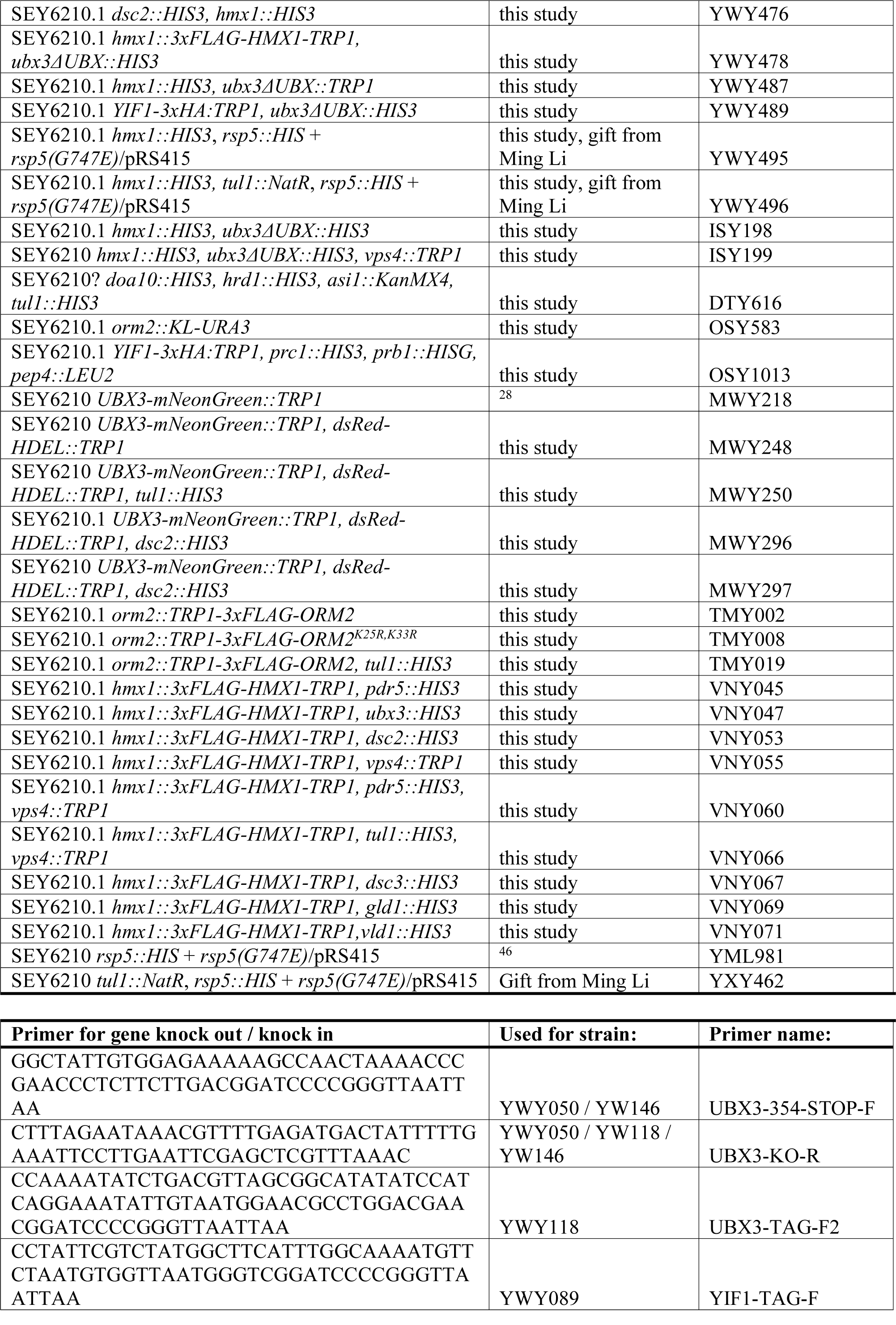
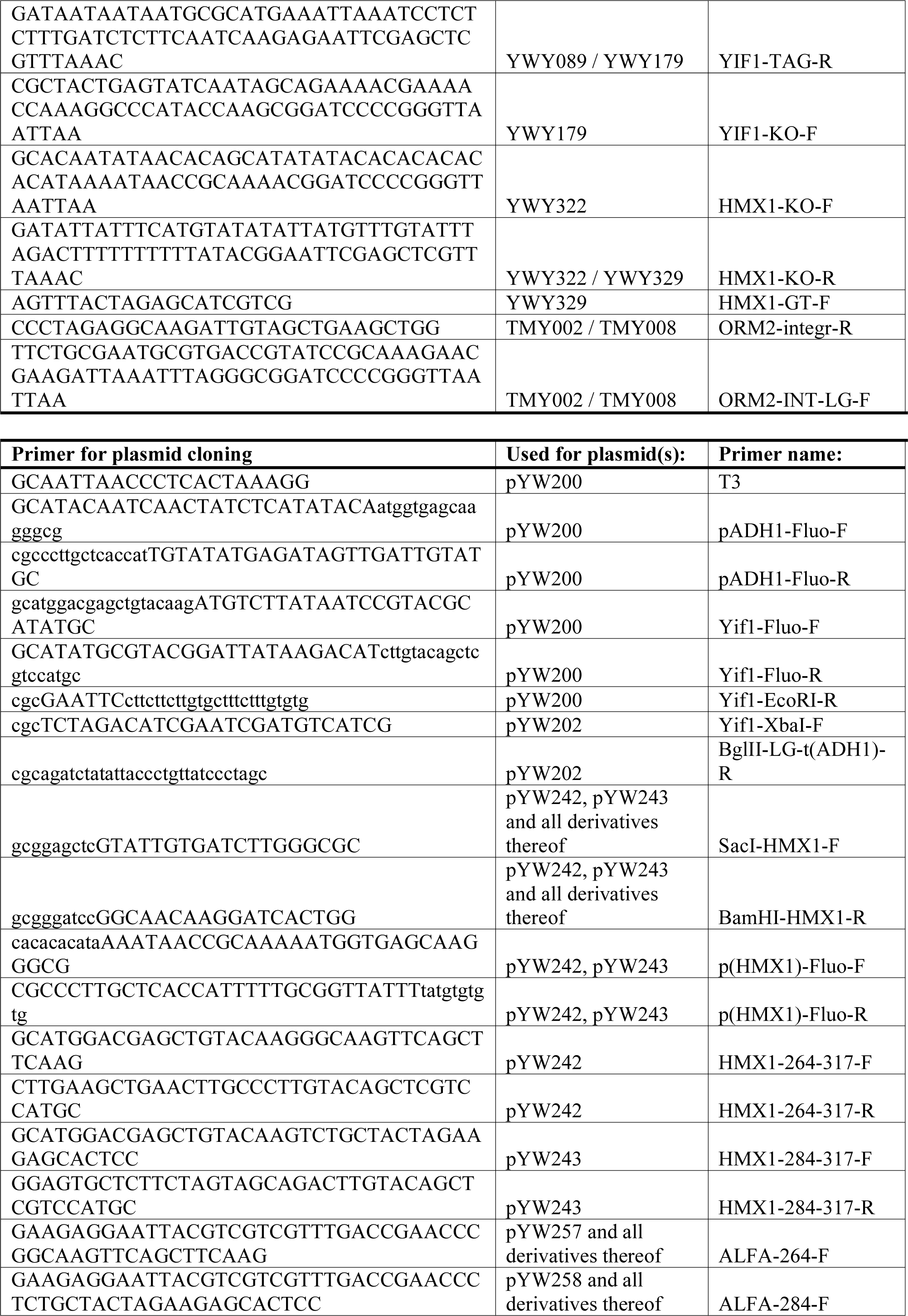
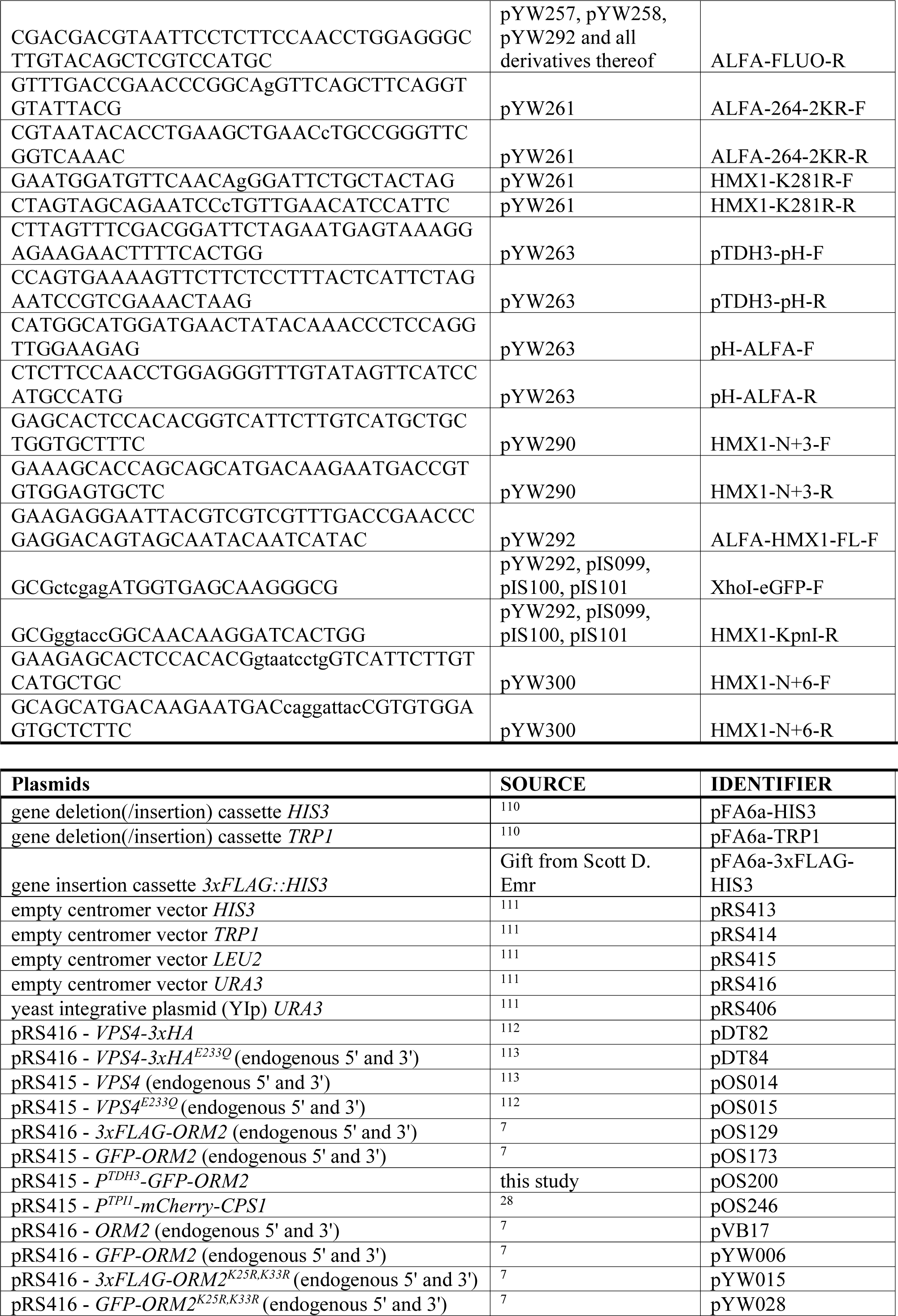
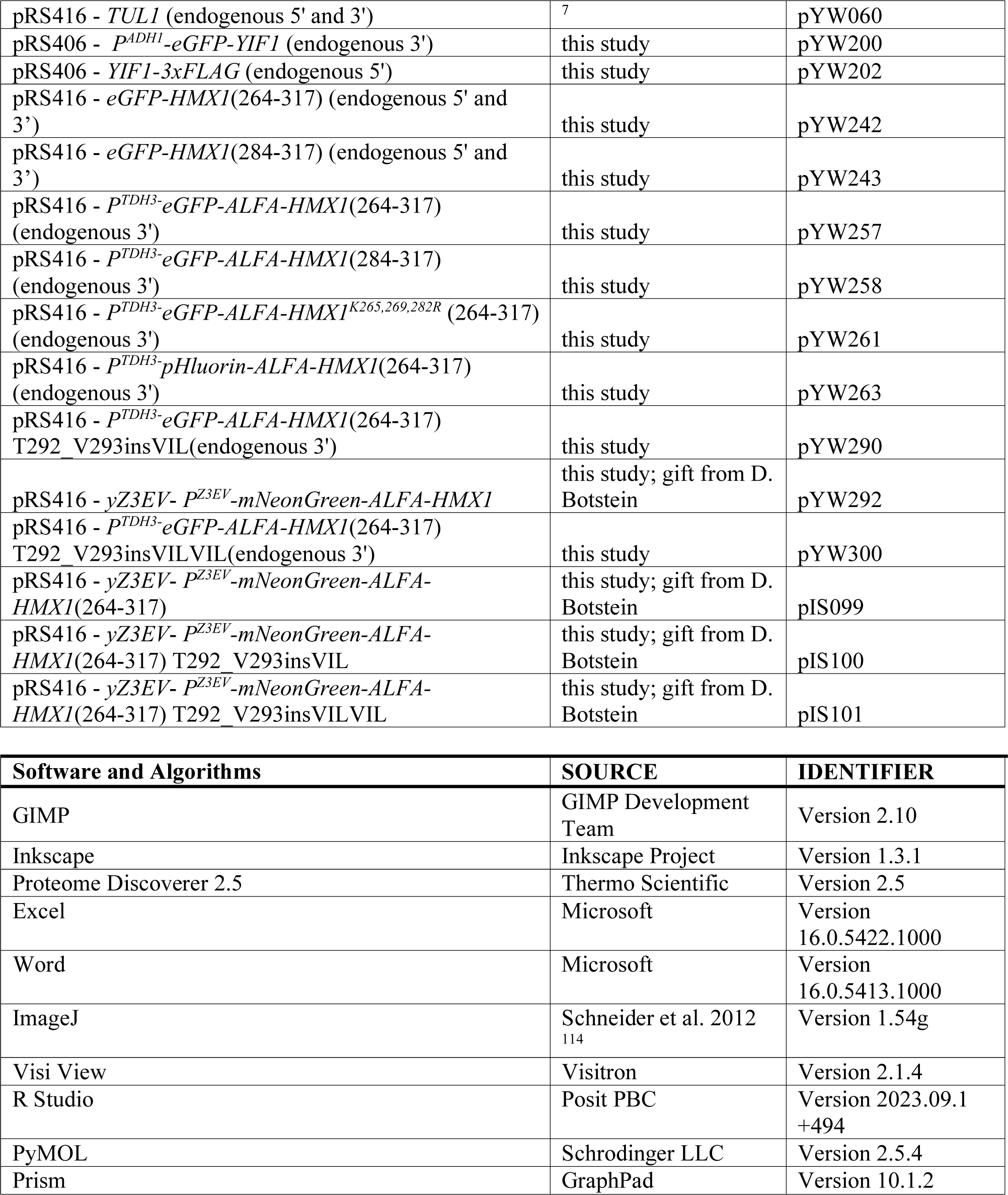
Reagents, strains and plasmid.

## Material and methods

### Yeast strains, plasmids, cloning, reagents and growth conditions

*S. cerevisiae* strains in this study were SEY6210 derivatives. All yeast strains, plasmids and reagents are listed in table S6. For liquid cultures, cells were incubated in YNB synthetic medium supplemented with amino acids and 2% glucose at 26°C in a shaker and were grown to midlog phase (OD_600_=0.5 – 1) for all experiments. Temperature sensitive mutants were shifted to restrictive temperature (37°C or 30°C) as indicated in the detailed methods section.

Genetic modifications were performed by PCR and/or homologous recombination using standard techniques. Plasmid-expressed genes including their native promoters and terminators were amplified from yeast genomic DNA and cloned into centromeric vectors (pRS series). All constructs were analyzed by DNA-sequencing and transformed into yeast cells using standard techniques. ß-Estradiol inducible plasmids are based on RB3579 and are derivatives of non-inducible pRS series plasmids. In detail, the entire ORF of yeGFP was excised with XhoI and KpnI, and replaced with the desired ORF of the gene of interest, which was flanked by XhoI and KpnI restriction sites. The desired ORF was amplified from existing plasmids with a forward primer that contained the starting methionine and the downstream 3’ region of M1, and a reverse primer that annealed within the 3’UTR of the respective gene of interest. Endogenously FLAG-tagged copies of *ORM2/HMX1* were cloned into pFA6a-*TRP1* plasmids and amplified by PCR with primers homologous to the respective promoter and 3’UTR of the gene of interest. This generated a cassette containing the gene of interest with the *TRP1* selection marker. This product was then integrated by homologous recombination into SEY6210.1 strains containing either a disrupted *orm2::URA3* or *hmx1::HIS3* locus. Integrants were counter selected and verified with DNA-sequencing. All yeast strains and plasmids used in this study as well as primer for PCR-based genetic modifications and cloning are listed table S6. pRS416-yZ3EV-Z3pr-yEGFP (RB3579) was a gift from David Botstein (Addgene plasmid # 69100; RRID:Addgene_69100)

### SILAC labeling and sample preparation for quantitative mass spectrometry

For quantitative analysis of protein abundances *tul1Δ* and *vps4Δ* mutants and their respective plasmid rescued WT cells were grown in synthetic medium (YNB) containing 2% glucose with all required amino acids except for uracil and lysine. Cells were pre-cultured for 24 hours in 10 mL medium at 26 °C. WT cells were grown with “heavy” [^13^C_6_^15^N_2_]-L-lysine (Sigma) at a final concentration of 20 mg/L and the *tul1Δ* and *vps4Δ* mutant cells were grown with “light” [^12^C_6_^14^N_2_]-L-lysine at a final concentration of 20 mg/L. Cells were grown to mid-log phase (OD600 ≤ 0.8) for 12 generations (24hours). After two washes at 4 °C with cold YNB medium without amino acids, WT “heavy” cells were mixed with an equal number of “light” mutant cells, filtered over a nitrocellulose membrane with a pore size of 0.2 µm (Amersham) and frozen in liquid nitrogen. Frozen cells were ground in a 6770 freezer mill (SPEX Sample Prep) with liquid nitrogen cooling and ground material was stored at -80 °C.

An aliquot of 1 mg dried cell lysate was dissolved in 200 µL ammonium hydrogen carbonate buffer (100 mM, pH 8) containing 0.5 % (w/w) sodium deoxycholate (SDO). The proteins were reduced by adding 20 µL of 100 mM DTT (30 min at 56 °C), alkylated with 20 µL 550 mM IAA (20 min at RT), and quenched with 20 µL 1 M DTT. Half of the sample was digested with 10 µg trypsin (Promega, 18 h at 37 °C). DOC was precipitated with 10 µL concentrated formic acid (conc. FA) and centrifuged at 36,000 × g for 10 min (supernatant 1). The pellet was redissolved in 40 µL 1 M ammonium solution, reprecipitated with 7 µL conc. FA and centrifuged at 36,000 × g for 10 min (supernatant 2). The two supernatants were combined and subjected to high pH fractionation using Pierce High pH Reversed-Phase Peptide Fractionation Kit (Thermo Scientific), according to the manufacturer’s instructions.

### LC-MS/MS analysis

High pH fractions were analyzed using an UltiMate 3000 nano-HPLC system coupled to a Orbitrap Eclipse mass spectrometer (Thermo Scientific, Bremen, Germany). The peptides were separated on a homemade fritless fused-silica microcapillary column (100 µm i.d. x 280 µm o.d. x 16 cm length) packed with 2,4 µm reversed-phase C18 material (Reprosil). Solvents for HPLC were 0.1% formic acid (solvent A) and 0.1% formic acid in 85% acetonitrile (solvent B). The gradient profile was as follows: 0-4 min, 4% B; 4-117 min, 4-30% B; 117-122 min, 30-100% B, and 122-127 min, 100 % B. The flow rate was 300 nL/min.

The Orbitrap Eclipse mass spectrometer equipped with a field asymmetric ion mobility spectrometer (FAIMS) interface was operating in the data dependent mode with compensation voltages (CV) of -45, -55 and -75 and a cycle time of one second. Survey full scan MS spectra were acquired from 375 to 1500 m/z at a resolution of 120,000 with an isolation window of 1.2 mass-to-charge ratio (m/z), a maximum injection time (IT) of 50 ms, and automatic gain control (AGC) target 400 000. The MS2 spectra were measured in the Orbitrap analyzer at a resolution of 15,000 with a maximum IT of 22 ms, and AGC target or 150 000. The selected isotope patterns were fragmented by higher-energy collisional dissociation with normalized collision energy of 30%.

### Analysis of quantitative proteome data and TMD analysis

Data Analysis was performed using Proteome Discoverer 2.5 (Thermo Scientific) with search engine Sequest. The raw files were searched against yeast database (orf_trans_all.fasta). Precursor and fragment mass tolerance was set to 10 ppm and 0.02 Da, respectively, and up to two missed cleavages were allowed. Carbamidomethylation of cysteine was set as static modification, oxidation of methionine and heavy lysine was set as variable modifications. Peptide identifications were filtered at 1% false discovery rate.

Three independent biological replicates were measured and analyzed. For further analysis only proteins with two or more H/L peptides that were quantified in at least two of the three biological experiments were considered. Proteins with a -log_2_ H/L ratio ≥ 0.3 and a p-value ≥ 0.05 were considered as significantly upregulated and compared against the landscape of yeast membrane proteins ^31^. Proteins which fulfilled these criteria were submitted to the DeepTMHMM server ^34^, subsequently the amino acid sequence of the transmembrane domains of membrane proteins were isolated and aligned with respect to their bilayer orientation, from the cytosolic side to the luminal side, with a custom-made R script listed.

Gene ontology (GO) term analysis was performed using the upregulated transmembrane standard names. They were mapped against the generic component terms using the GoSLIM mapper of the *Saccharomyces* genome database ^91^. GO terms with only one hit were removed and hits from the term ‘cytoplasmic vesicles’ were fused either with the plasma membrane or with endosomes according to their localization in the yeast GFP database ^92^. The ratio of the observed (dataset frequency) vs. the expected number of genes (genome frequency) associated with the GO term was also calculated and referred to as enrichment over genome ^93^.

### Molecular dynamics simulation

We retrieved the Dsc2 starting structure from the AlphaFold Protein Structure Database (https://alphafold.ebi.ac.uk/, accession: AF Q08232 F1). We focused the analysis on the rhomboid like domain of Dsc2, and therefore omitted an β-sheet (aa 148 – 182) that connects TM3 with TM4 as well as the undefined C-terminal portion (aa 241 – 322). Omitting these regions also accelerate the convergence of the simulation. Preparation of the initial structure, comprising the protonation and minimization were conducted in the MOE platform (Molecular Operating Environment, version 2022.02, Molecular Computing Group Inc, Montreal, Canada). Dsc2 was oriented in the membrane using PPM 2.0 ^94^ and embedded in a lipid bilayer comprising 45% POPC, 45% POPE and 10% cholesterol with the CHARMM-GUI Membrane Builder ^95^. A 75 Å*75 Å membrane with a default 22.5 Å aqueous layer was added to ensure the efficient embedding of the Dsc2 and minimize artificial interference among neighboring images during the simulation. The resulting system was parameterized with LEaP from the AmberTools23 ^96^ using TIP3P ^97^, FF14SB ^98^ and Lipid21 ^99^ as force fields for water, protein and lipids. We equilibrated the system by minimizing gradually all hydrogen atoms, heavy atoms of water, lipids and the protein. The system was then heated up from 100K to 300K, minimized to cool down before being heated up again to 300K. Finally, we equilibrated the system with 1ns isotropic position scaling followed by 10ns pressure scaling, as was suggested by previous studies ^100,101^. The equilibrated system was simulated with periodic boundary conditions using the SHAKE algorithm for hydrogens under constant pressure for 100ns with amber 22 ^96^. For non-bonding interactions a 8 Å cut off was applied and long-range electrostatics were estimated with the particle mesh Ewald method ^102^. We took the last 40ns where the simulation converges to obtain statistics on membrane thickness changes with the membplugin ^103^ and visualized the dynamics with VMD ^104^.

### Preparation of whole cell protein extracts

To prepare whole cell lysates, proteins were extracted by a modified alkaline extraction protocol^105^. Briefly, 4 OD_600_ of logarithmically growing yeast were harvested by centrifugation (5 min, 4,000 rpm, 4 °C) and washed in ice cold 10 mM NaF. After centrifugation (3 min, 13,000 rpm, 4 °C) cells were resuspended in 0.1M NaOH and incubated at room temperature for 5 min. After centrifugation (3 min, 13,000 rpm, 4 °C) pellets were resuspended in Lämmli sample buffer (60 mM Tris-HCl pH 7.5, 2% SDS, 1% β-mercaptoethanol, 10% glycerol, bromophenol blue), denatured (95 °C, 5 min), and cell debris was removed by centrifugation (3 min, 13,000 rpm, 4 °C).

### Western Blot analysis and immuno-detection

Protein extracts dissolved in Lämmli sample buffer or SDS sample buffer were separated by SDS-PAGE (Biorad Mini Protean) and transferred to ethanol activated Amersham Hybond-P 0.45 µm PVDF membranes (Cytiva, Cat. #10600023) by either semi-dry or wet electro-blotting with ethanol based transfer buffer (25 mM Tris-HCl, 192 mM Glycine, 20% EtOH, 0.1% SDS). Membranes were blocked in 5% milk in TBS-T for 1 h at room temperature, and incubated with primary antibodies over night at 4 °C. Membranes were afterwards washed 5 – 10 times with TBS-T (50 mM Tris-HCl pH 7.5, 500 mM NaCl, 0.05% Tween-20). Secondary antibodies conjugated to HRP (horseradish peroxidase) were diluted in 5% Milk in TBS-T and added to the membranes for 60 – 90 min at room temperature, followed by 5 – 10 washing steps with TBS-T. Membranes were developed using ECL HRP substrate (Advansta, Cat. #541005X). Oxidised Luminol was detected using medical X-Ray films. Films were scanned and further processed with GIMP (version 2.10). No non-linear processing was applied. Antibodies used in this study are listed in table S6. The α-Dsc2, α-Dsc3, α-Tul1 and α-Ubx3 antibodies were kindly provided by Peter J. Espenshade, Johns Hopkins University School of Medicine.

### Immunodetection of ubiquitinated proteins

For detection of ubiquitinated proteins, samples were separated by SDS-PAGE gel electrophoresis using 12.5% gels. Proteins were then transferred to Amersham Hybond-P 0.45 µm PVDF membranes (Cytiva, Cat. #10600023) by wet blotting (80 V, 2 h) at 4 °C with ethanol based transfer buffer (25 mM Tris-HCl, 192 mM Glycine, 20% EtOH, 0.1% SDS). After transfer, the membrane was blocked with 5% BSA in TBS-T (50 mM Tris-HCl pH 7.5, 150 mM NaCl, 0.45% Tween-20) before overnight incubation with P4D1 anti-ubiquitin antibody (Santa Cruz, Cat. # sc-8017) with 0.45% Tween-20 in 5% BSA blocking solution. Washing was done as described above. Membranes were developed using ECL select (Cytiva, Cat. # GERPN2235).

### Denaturing immunoprecipitation of ubiquitinated species

Denaturing immunoprecipitation of FLAG-Hmx1 and truncated versions of eGFP-ALFA-Hmx1 was adapted from (Schmidt et al., 2019). Briefly, 50 OD_600_ (150 OD_600_ for downstream ubiquitin linkage immunoprecipitation) of logarithmically growing yeast were collected with 1 mL ice-cold 10 mM NaF. Cell pellets were resuspended in 0.1M NaOH and incubated for 5 min at RT. After centrifugation (3 min, 13,000 rpm, 4 °C), pellets were resuspended in 200 µL SDS lysis buffer (50 mM Tris-HCl pH 8, 1 mM EDTA, 1% SDS, 2 M urea, 10 mM NaF) supplemented with protease inhibitors (1x Complete® EDTA-free (Roche, Cat. # C7698), 1 mM PMSF, 10 mM N-ethyl maleinimid). Pellets were then solubilized by vortexing with glass beads (0.75-1 mm; 10 min at RT) and denatured (42 °C, 1 h). Lysates were diluted by addition of 1800 µL immunoprecipitation (IP) buffer (50 mM Tris-HCl pH 8, 100 mM NaCl, 1% Triton X-100, supplemented with protease inhibitors including yeast protease inhibitors (Sigma, Cat. # P8215) and cleared by centrifugation (10 min; 13,000g; 4 °C). Supernatants were then added to prewashed anti-FLAG M2 magnetic beads (Sigma, Cat. # M8823) or anti-ALFA ST magnetic beads (NanoTag Biotechnologies, Cat. # N1516) and immunoprecipitated for 2 – 3 h at 4 °C. The beads were washed twice with IP buffer (50 mM Tris-HCl, pH 8, 100 mM NaCl, 1% Triton X-100) and thrice with wash buffer (50 mM Tris-HCl pH 8, 300 mM NaCl, 0.1% Tween-20) for 5 min at 4 °C. FLAG-tagged proteins were eluted by addition of 40 µL IP buffer supplemented with 500 ng/mL 3xFLAG peptide (Sigma, Cat. # F4799) and incubated for 1 h at room temperature. Magnetic beads were removed and the eluate was supplemented with 5x SDS sample buffer (250 mM Tris-HCl pH 6.8, 10% SDS, 50% glycerol, 10% β-mercaptoethanol) and denatured for 10 min at 95 °C. ALFA-tagged proteins were eluted with 50 µL 2x SDS sample buffer (100 mM Tris-HCl pH 6.8, 4% SDS, 20% glycerol, 4% β-mercaptoethanol) and denatured for 10 min at 95 °C.

For further immunoprecipitation with K48 or K63 selector resins, FLAG-proteins were eluted with 120 µL IP buffer as described above. ALFA-tagged proteins were immunoprecipitated with anti-ALFA CE agarose beads (NanoTag Biotechnologies, Cat. # N1512-L) instead of anti-ALFA ST and eluted twice with 60 µL IP buffer supplemented with 800 µg/mL ALFA peptide (NanoTag Biotechnologies) and incubated for 15 min at 37 °C. In both cases, eluates were split into three parts. The fraction for detection of holo-ubiquitinated species was denatured immediately after elution with 5x SDS sample buffer for 10 min at 95 °C. The remaining fraction was diluted with 720 µL IP buffer supplemented with protease inhibitors (1x Complete® EDTA-free (Roche, Cat. # C7698), 1 mM PMSF, 10 mM N-ethyl maleinimid, yeast protease inhibitors (Sigma, Cat. # P8215)) and equally split to anti-Ubiquitin K48 Selector agarose beads (NanoTag Biotechnologies, Cat. # N1810) and anti-Ubiquitin K63 Selector agarose beads (NanoTag Biotechnologies, Cat. # N1910). Beads were incubated for 1 h at 4 °C followed by two washes with IP buffer and three washes with Wash buffer. Ubiquitinated species were eluted and denatured with 2x SDS sample buffer for 10 min at 95 °C.

### Non-denaturing immunoprecipitation

Non-denaturing isolation of FLAG-Orm2 and Ubx3-FLAG was performed essentially as described before (Schmidt et al., 2019) from 1000 OD_600_ of cryo-ground cells. For non-denaturing isolation of FLAG-Hmx1, for truncated versions of eGFP-ALFA-Hmx1 and Yif1-HA, logarithmically growing cells (100 OD_600_ for Hmx1, 50 OD_600_ for HA) were harvested by centrifugation with ice cold 10 mM NaF, and pellets were frozen in liquid nitrogen. Frozen pellets were resuspended in 500 µL lysis buffer (50 mM HEPES/KOH, pH 6.8; 150 mM KOAc, 2mM Mg(OAc)_2_; 1mM CaCl_2_; 15% glycerol) supplemented with protease inhibitors (1x Complete® EDTA-free (Roche, Cat. # C7698), 1 mM PMSF, 10 mM N-ethyl maleinimid, yeast protease inhibitors (Sigma, Cat. # P8215)) and solubilized by vortexing with glass beads (0.75-1 mm; 10 min at 4 °C). Lysates were then diluted with 500 µL lysis buffer supplemented with protease inhibitors and 2% glyco-diosgenin (Sigma, Cat. # 850525) and incubated rotating at 4 °C for 1h. Unsolubilized material was removed by centrifugation (13,000 × g; 15 min; 4 °C). The supernatant (800 µL) was added to equilibrated anti-FLAG M2 magnetic beads (Sigma, Cat. # M8823), anti-ALFA ST magnetic beads (NanoTag Biotechnologies, Cat. # N1516) or anti-HA magnetic beads (Thermo Fisher, Cat. #88836) and incubated for 2 – 3 h at 4 °C. Beads were recovered with a magnetic rack and washed two times with IP buffer containing 0.1% glyco-diosgenin and three times with IP buffer containing 0.1% glyco-diosgenin and 300 mM NaCl for 5 minutes at 4 °C. FLAG-proteins were eluted with 50 µL 500 ng/mL 3xFLAG peptide (Sigma, Cat. # F4799) at RT for 1h and supplemented with 5x SDS sample buffer (250 mM Tris-HCl pH 6.8, 10% SDS, 50% glycerol, 10% β-mercaptoethanol) and denatured for 10 min at 95 °C. ALFA-tagged and HA-tagged proteins were eluted with 50 µL 2x SDS sample buffer (100 mM Tris-HCl pH 6.8, 4% SDS, 20% glycerol, 4% β-mercaptoethanol) and denatured for 10 min at 95 °C.

### Size Exclusion Chromatography High Performance Liquid Chromatography (SEC-HPLC)

Non denaturing immunoprecipitation of Ubx3-FLAG was performed as described above. Prior to elution, beads were additionally washed two times with PBS supplemented with 0.1% glyco-diosgenin (Sigma, Cat. # 850525). Bound material was eluted (1h at RT, shaking at 850 rpm) from anti-FLAG M2 magnetic beads (Sigma, Cat. # M8823) with homemade 10 µg/µL Strep-6xFLAG-peptide in PBS supplemented with 0.1% glyco-diosgenin. After elution, FLAG-beads were separated with a magnetic rack and PBS-washed MagStrep“type3“ XT Beads (IBA, Cat. #2-4090-002) were added to eliminate excess of Strep-6xFLAG-Peptide (1h at RT, shaking at 850 rpm). The supernatant of this purification were used for size exclusion chromatography (SEC).

The SEC analysis was performed utilizing an ÄKTA Ettan LC system coupled with a Superose 6 Increase 3.2/300 column. The column was pre-equilibrated with a mobile phase composed of PBS with 0.05% GDN (25x critical micelle concentration (CMC)). Calibration was executed using molecular weight standards, including Thyroglobulin (669 kDa), Ferritin (440 kDa), Aldolase (158 kDa), Ovalbumin (44 kDa), and Aprotinin (6.5 kDa). For sample analysis, 100 μl of eluate was loaded onto the ÄKTA Ettan LC system, employing the previously described mobile phase at a flow rate of 0.034 mL/min. Fraction collection was conducted in 80 μL aliquots using a fraction collector. After collection, fractions were analyzed by SDS PAGE and Western Blot analysis.

### Live cell fluorescence Microscopy

Live cell epifluorescence microscopy was carried out using a Zeiss Axio Imager M1 equipped with a SPOT Xplorer CCD camera, standard fluorescent filters and Visitron VisiView software (version 2.1.4). For microscopy, cells were grown to midlog (OD_600_ 0.5 – 0.8) phase in YNB media, concentrated by centrifugation, resuspended in A.d. and directly mounted onto glass slides. Brightness and contrast of the images in the figure were adjusted using ImageJ software (version 1.54g).

### Cycloheximide chase assay

Logarithmically growing cells (20 OD_600_) were harvested by centrifugation. Cells were resuspended in 50 mL fresh medium (for experiments with temperature-sensitive mutants the medium was pre-warmed at 30 °C). For chemical proteasome inhibition *pdr5Δ* cells were pre-incubated with 50 µM MG-132 (Sigma, Cat. # 474787) for 10 minutes. At t = 0 min 10 mL (4 OD_600_) were harvested by centrifugation, washed once with ice-cold 10 mM NaF (see above), and pellets were snap frozen in liquid nitrogen. To the remaining culture 50 µg/mL cycloheximide (Sigma, Cat. # C7698) was added from a 10 mg/mL stock. After the indicated time points 10 mL culture were harvested, washed and frozen as above. Whole cell extracts were prepared by alkaline extraction. SDS-PAGE, Western blot detection and quantification were done as described above. For experiments involving proteasome inhibition, strains harbor an additional *PDR5* deletion and were incubated with 50 µM or vehicle (DMSO) 10 min prior to the addition of CHX.

### Subcellular fractionation

The subcellular fractionation protocol was adapted from ^7,47,48^. Briefly, logarithmically growing *vps4Δ* CDC48 or cdc48-3 cells were shifted to 37 °C for 2 h and simultaneously treated with 50 µM MG-132 (Sigma, Cat. # 474787) or the respective vehicle. 40 OD_600_ of midlog phase (OD600=0.5 – 1) growing cells were collected and resuspended in 1 mL ice-cold 10 mM NaF. Cell pellets were resuspended in 500 µL MF Buffer (300 mM Sorbitol, 1 mM EDTA, 20 mM Tris-HCl pH 7.5, 100 mM NaCl), supplemented with protease inhibitors (1x Complete® EDTA-free (Roche, Cat. # C7698), 1 mM PMSF, 10 mM N-ethyl maleinimid, yeast protease inhibitors (Sigma, Cat. # P8215)), 100 µL 0.75 – 1 mm glass beads were added and vortexed (6 x 1 min at 4 °C). Cell lysates were cleared at 2500 × g for 5 min. 400 µL of the supernatant was ultracentrifuged at 100,000 × g for 1 – 1.5 h to separate pellet (P100) and supernatant (S100) fractions. 300 µL of the S100 fraction were removed diluted with IP buffer (50 mM Tris-HCl pH 8, 100 mM NaCl, 1% Triton X-100) supplemented with protease inhibitors (1x Complete® EDTA-free (Roche, Cat. # C7698), 1 mM PMSF, 10 mM N-ethyl maleinimid, yeast protease inhibitors (Sigma, Cat. # P8215) and added directly to the anti-FLAG M2 magnetic beads (Sigma, Cat. # M8823). The P100 was washed once in cold MF Buffer without disturbing the pellet and centrifuged at 13,000 × g for 5 min. The supernatant was discarded and the washed P100 fraction was resuspended in 200 µL SDS lysis buffer (50 mM Tris-HCl pH 8, 1 mM EDTA, 1% SDS, 2M urea, 10 mM NaF) supplemented with protease inhibitors (1x Complete® EDTA-free (Roche, Cat. # C7698), 1 mM PMSF, 10 mM N-ethyl maleinimid). An equivalent amount of P100 (compared to S100) was transferred to IP buffer (50 mM Tris-HCl pH 8, 100 mM NaCl, 1.0% Triton X-100) supplemented with protease inhibitors (1x Complete® EDTA-free (Roche, Cat. # C7698), 1 mM PMSF, 10 mM N-ethyl maleinimid, yeast protease inhibitors (Sigma, Cat. # P8215)) and added to the anti-FLAG M2 magnetic beads (Sigma, Cat. # M8823). Immunoprecipitations under denaturing conditions were described above.

### Full lipidome analysis

Isogenic WT and mutant strains were grown into logarithmic phase in minimal selection medium. Approximately 30 OD_600_ cells were rapidly harvested and frozen in liquid nitrogen. Pellets were resuspended in 1.5 mL sterile, ice-cold water and the OD_600_ was measured. Exactly 20 OD cells were transferred into a fresh tube and water added to a final volume of 1 mL. 400 µL 0.5 mm glass beads were added, and glass bead lysis was performed for 15 min at 4 °C. 750 µL of the lysate were transferred to fresh tubes, frozen in liquid nitrogen, and shipped to Lipotype (Dresden, Germany) for subsequent lipid extraction and shotgun mass spectrometric lipidomics analysis. Lipidomic profiling was conducted across four independent experiments. Quantitative data is usually displayed as mean ± standard deviation from at least 3 biological replicates. Statistical significance was tested in Prism software (version 10.1.2) by multiple t-tests correcting for multiple comparisons ^106^, with a false discovery rate Q = 1%, without assuming individual variance for each sample. P-values were flagged with > 0.05 (ns), ≤ 0.05 (*), ≤ 0.01 (**) and ≤ 0.001 (***).

### ß-Estradiol induced protein expression

To induce protein expression, 500 nM β-Estradiol (Sigma, Cat. #E2758) was added to logarithmically grown overnight cell cultures. Protein induction and/or localization was examined at the indicated time points using Western blot analysis and/or live cell fluorescent microscopy. To examine protein degradation kinetics, 4 OD_600_ of cells were harvested after 90 minutes of β-Estradiol induction (t=0) by centrifugation and snap frozen in liquid nitrogen. To the remaining culture 50 µg/mL cycloheximide (Sigma, Cat. # C7698) was added. At the indicated time points 4 OD_600_ of culture were harvested, and snap frozen in liquid nitrogen. Whole cell extracts were prepared by alkaline extraction. SDS-PAGE, Western blot detection, and quantification were done as described above.

### Quantification and line scans of Western Blot analysis

Western blot signals were quantified by densitometry using ImageJ software (version 1.54g). Quantifications were exported to Microsoft Excel (Version 16.0.5422.1000), normalized to the respective Pgk1 loading controls, and presented as mean ± standard deviation of the mean from at least three independent experiments. t = 0 was set to 1. Line scans of the S100 fraction (Figure 11D, E) were measured using the plot profile function of ImageJ software (version 1.54g). Briefly, a 25.4 mm long line was drawn and placed at three different vertical positions per each lane. The numerical values were extracted from the resulting plots transferred to Microsoft Excel (Version 16.0.5422.1000), and the values were inverted by subtraction from 255. The mean values, the standard deviations and the 95% confidence intervals were calculated for each lane and visualized with R Studio (version 2023.09.1 +494).

## Data and software availability

All relevant data has been included in the paper in main figures and supplemental information.

