## Supplementary figures and images for "The Dsc ubiquitin ligase complex identifies transmembrane degrons to degrade orphaned proteins at the Golgi"

### Figure S1-S7

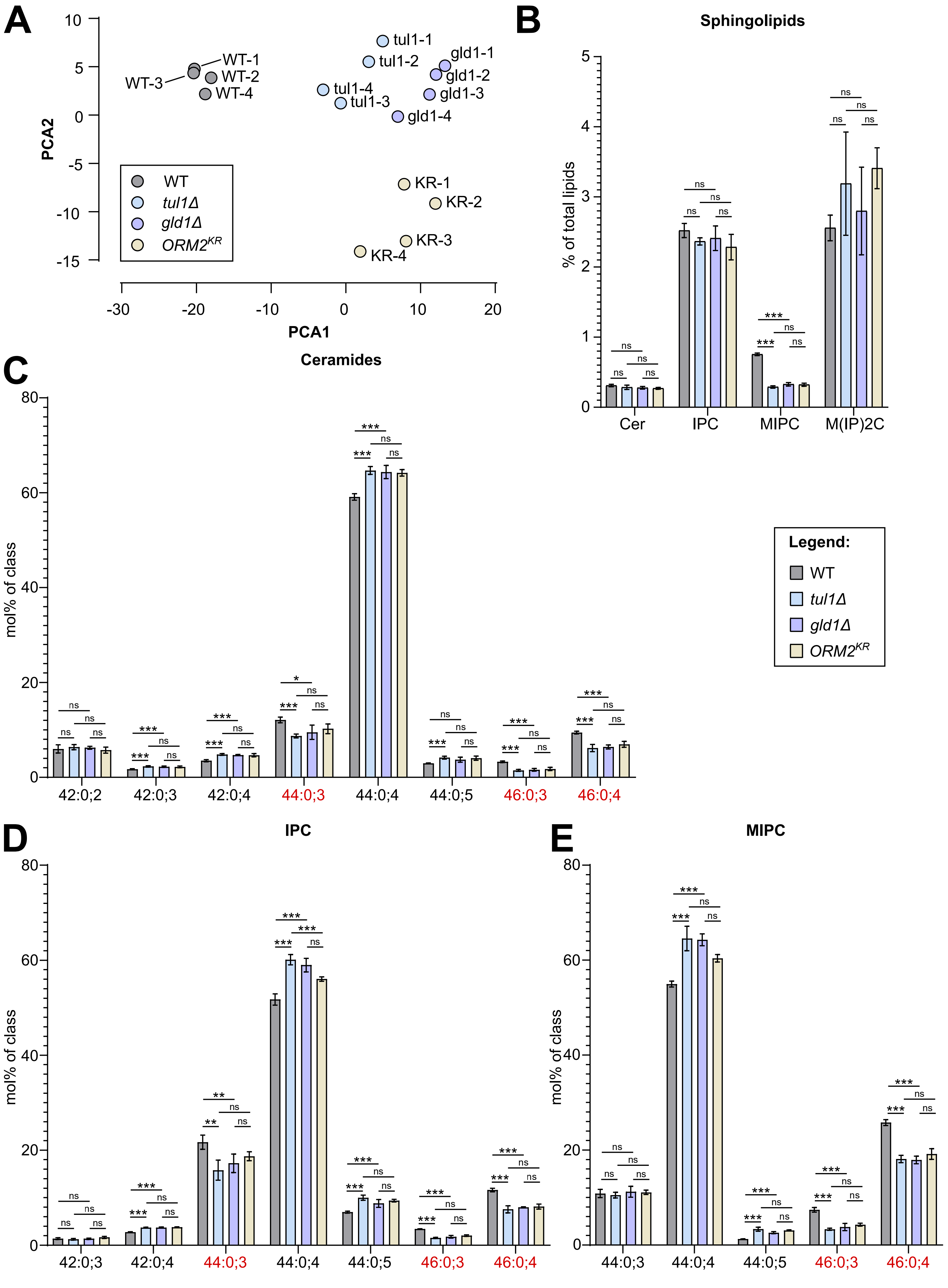

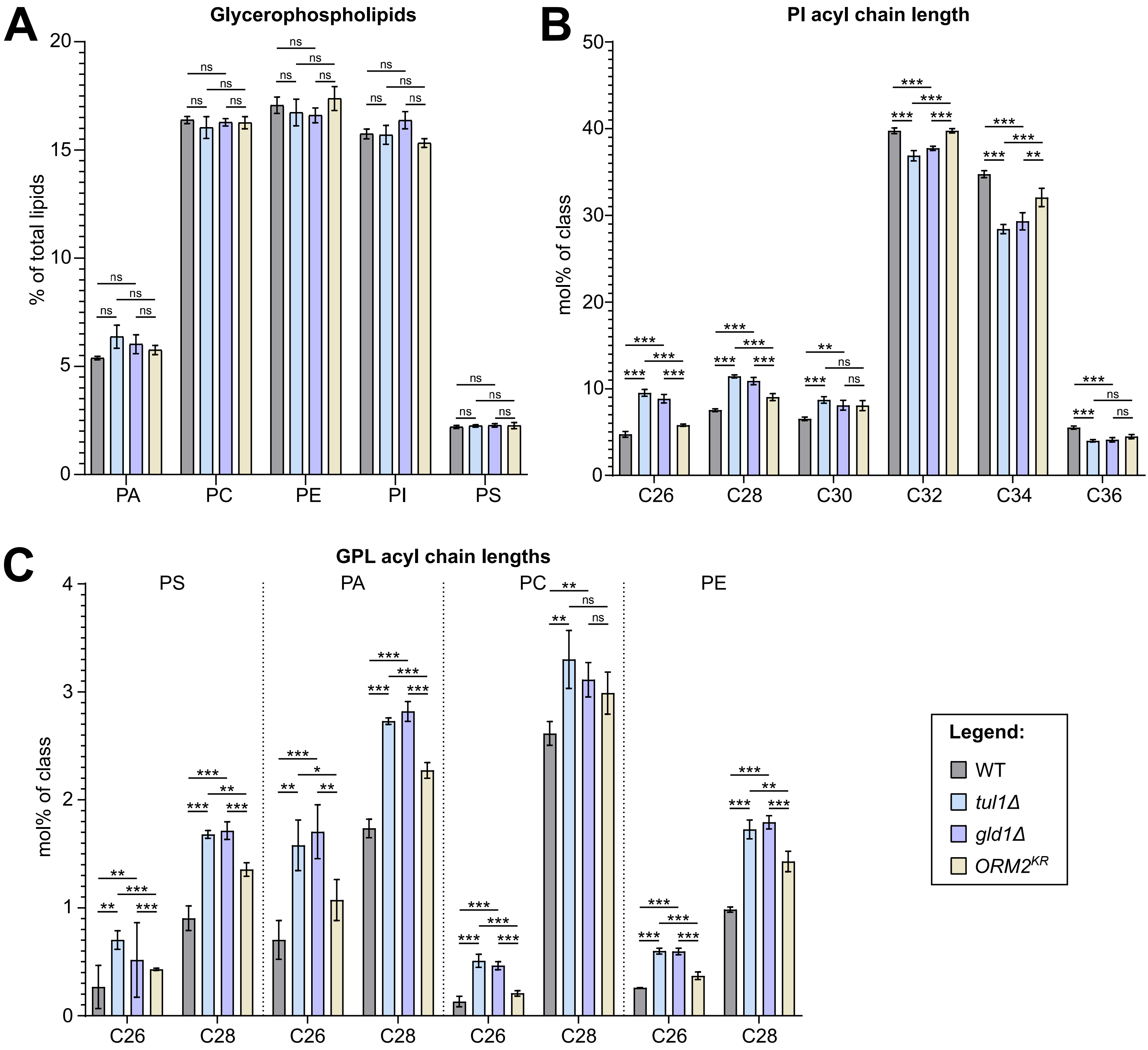

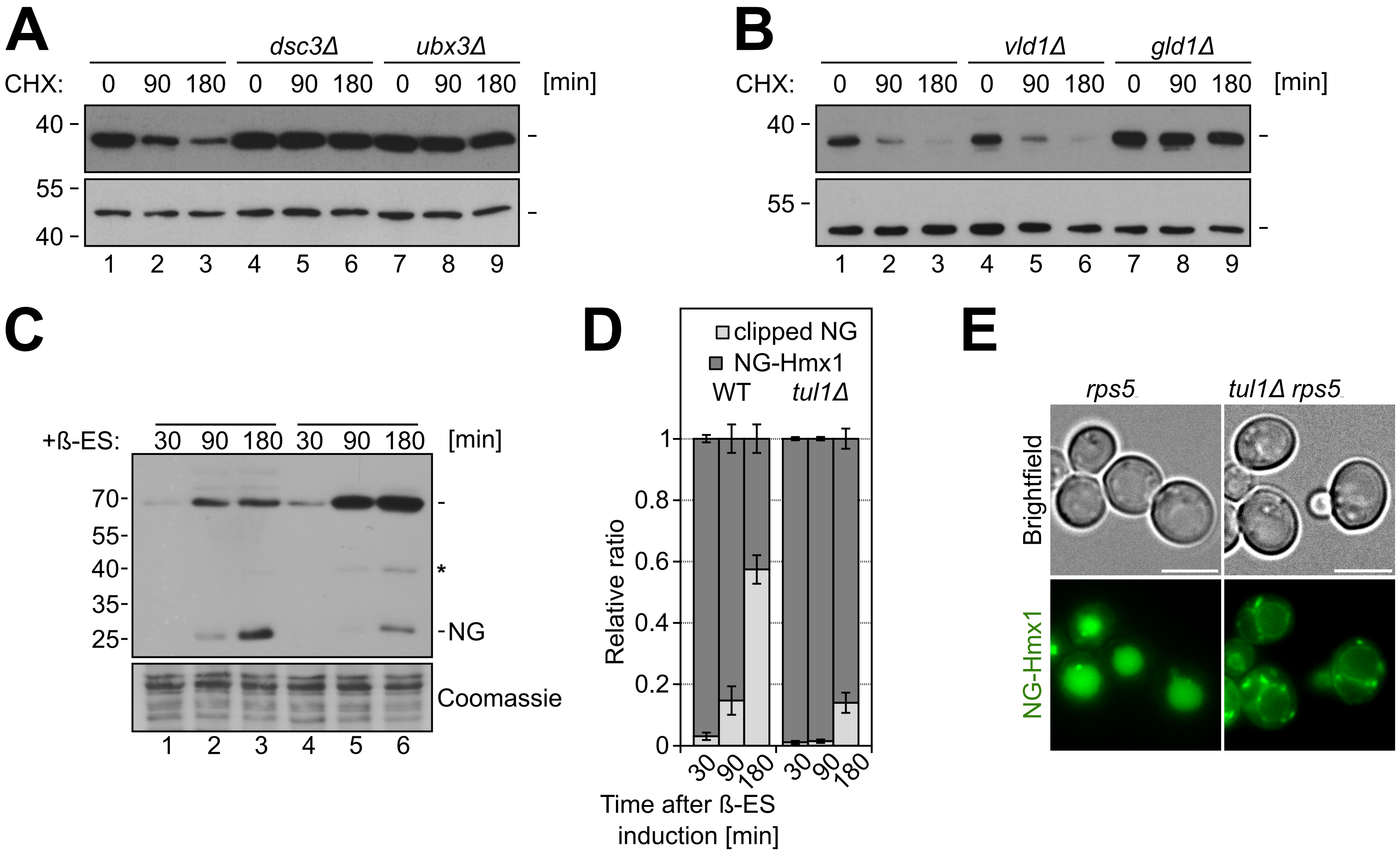

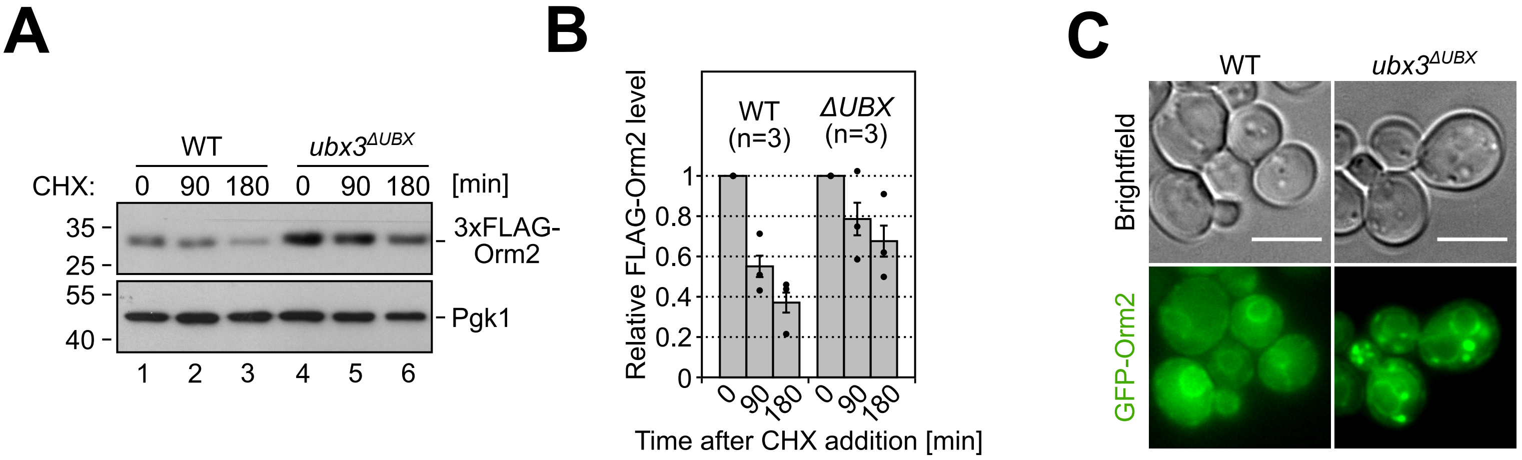

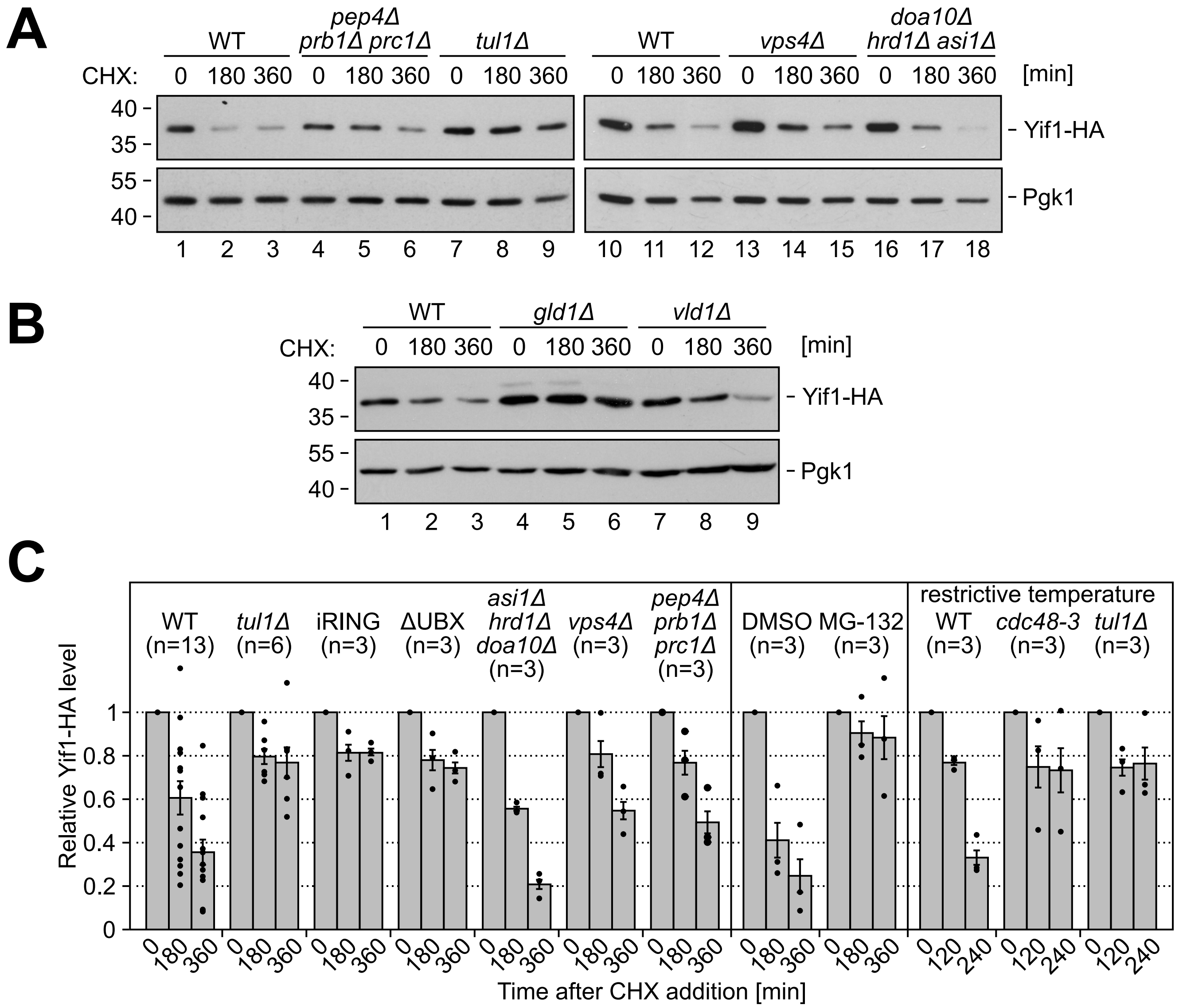

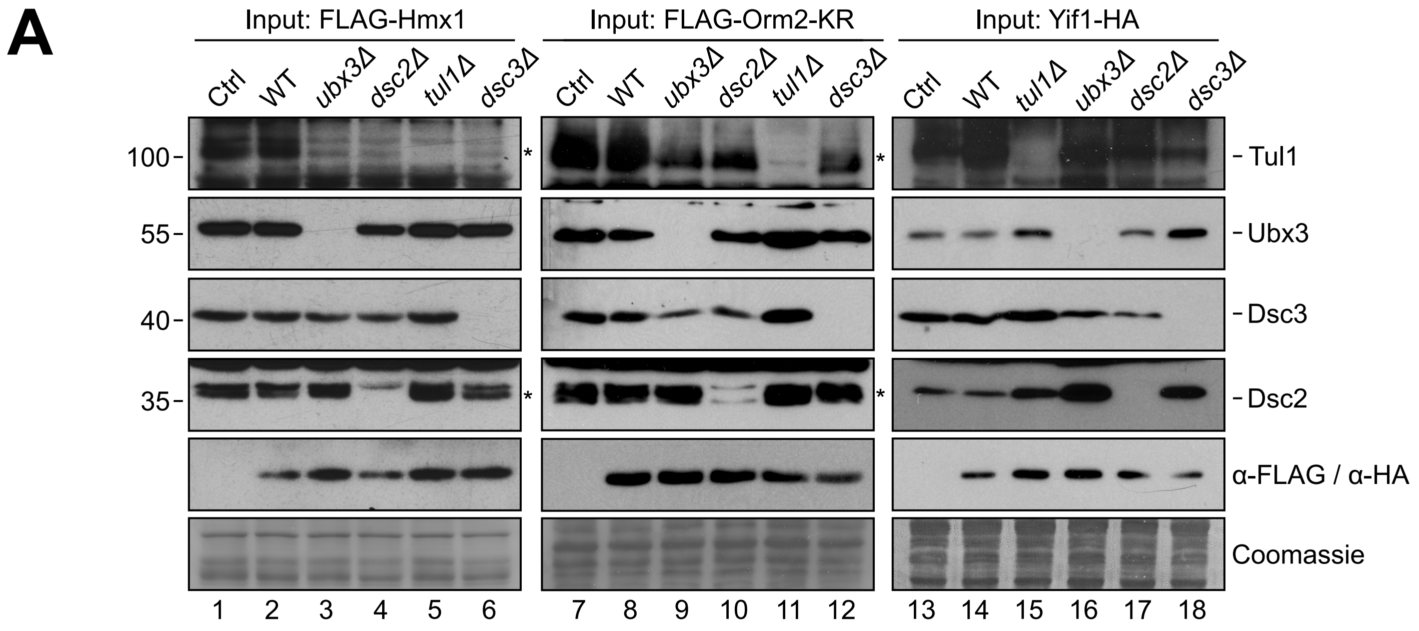

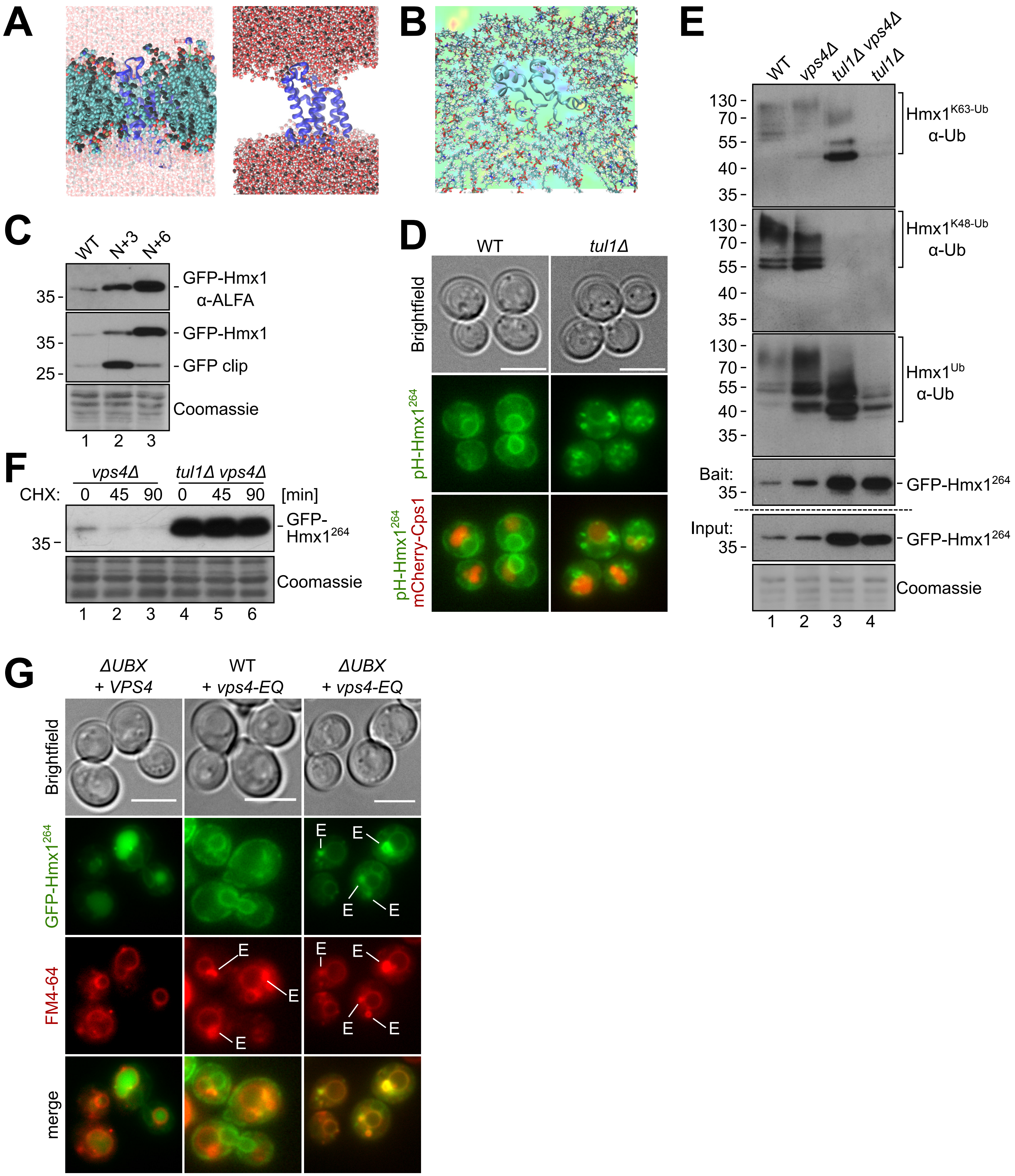
